# Heat Shock Factor 1 (HSF1) as a new tethering factor for ESR1 supporting its action in breast cancer

**DOI:** 10.1101/2021.05.06.442900

**Authors:** Natalia Vydra, Patryk Janus, Paweł Kuś, Tomasz Stokowy, Katarzyna Mrowiec, Agnieszka Toma-Jonik, Aleksandra Krzywon, Alexander Jorge Cortez, Bartosz Wojtaś, Bartłomiej Gielniewski, Roman Jaksik, Marek Kimmel, Wiesława Widłak

**Author notes:** These authors contributed equally (Natalia Vydra, Patryk Janus, Paweł Kuś). Correspondence (WW); (NV).

## Abstract

Heat shock factor 1 (HSF1), a key regulator of transcriptional responses to proteotoxic stress, was recently linked to estrogen (E2) signaling through ESR1. We found that an HSF1 deficiency could lead to the inhibition of the mitogenic action of E2 in breast cancer cells. The stimulatory effect of E2 on the transcriptome is weaker in HSF1-deficient cells, in part due to the higher basal expression of E2-dependent genes, which correlates with the enhanced binding of unliganded ESR1 to chromatin. HSF1 and ESR1 can cooperate directly in E2-stimulated regulation of transcription, and HSF1 potentiates the action of ESR1 through a mechanism involving chromatin reorganization. Analyses of data from the TCGA database indicate that HSF1 increases the transcriptome diversity in ER-positive breast cancer and can enhance the genomic action of ESR1. Moreover, *ESR1* and *HSF1* are only prognostic when analyzed together (the worst prognosis for ER−/HSF1^high^ cancers).

## Introduction

Breast cancer is the most common malignancy in women worldwide. Four clinically relevant molecular types are distinguished based on the expression of estrogen receptors (ERs) and HER2 (ERBB2). Among them, luminal adenocarcinomas, characterized by the expression of estrogen receptors, constitute about 70% of all breast cancer cases. There are two classical nuclear estrogen receptors, ERα (ESR1) and ERβ (ESR2), and structurally different GPR30 (GPER1), which is a member of the rhodopsin-like family of the G protein-coupled and seven-transmembrane receptors. ERα expression is most common in breast cancer and its evaluation is the basis for determining the ER status. Activity of estrogen receptors is modulated by steroid hormones, mainly estrogens, which are synthesized from cholesterol. According to epidemiological and experimental data, estrogens alongside the mutations in *BRCA1* and *BRCA2*, *CHEK2*, *TP53*, *STK11* (*LKB1*), *PIK3CA*, *PTEN*, and other genes, are key etiological factors of breast cancer development (Yaşar et al., 2017) (Verigos and Magklara, 2015). The mechanism of estrogen-stimulated breast carcinogenesis is not clear. According to the widely accepted hypothesis, estrogens acting through ERα, stimulate cell proliferation and can support the growth of cells harboring mutations which then accumulate, ultimately resulting in cancer. Another hypothesis suggests the ERα-independent action of estrogens via their metabolites, which can exert genotoxic effects, contributing to cancer development (Yager and Davidson, 2006) (Pescatori et al., 2021).

Previously, we have found that the major female sex hormone 17β-estradiol (E2) stimulates activation of the Heat Shock Factor 1 (HSF1) in estrogen-dependent breast cancer cells via MAPK signaling (Vydra et al., 2019). HSF1 is a well-known regulator of cellular stress response induced by various environmental stimuli. It mainly regulates the expression of the Heat Shock Proteins (HSPs), which function as molecular chaperones and regulate protein homeostasis (Ran et al., 2007). HSF1-regulated chaperones control, among other, the activity of estrogen receptors (Echeverria and Picard, 2010). ERs remain in an inactive state trapped in multimolecular chaperone complexes organized around HSP90, containing p23 (PTGES3), and immunophilins (FKBP4 or FKPB5) (Segnitz and Gehring, 1995). Upon binding to E2, ERs dissociate from the chaperone complexes and become competent to dimerize and regulate the transcription. ERs bind DNA directly, to the estrogen-response elements, EREs, or act indirectly through recruiting transcriptional co-activators (Heldring et al., 2007) (Renoir, 2012). HSP90 is essential for ERα hormone binding (Fliss et al., 2000), dimer formation (Powell et al., 2010), and binding to the EREs (Inano et al., 1994). Also, the passage of the ER to the cell membrane requires association with the HSP27 (HSPB1) oligomers in the cytoplasm (Razandi et al., 2010). More than 20 chaperones and co-chaperones associated with ERα in human cells have been identified through a quantitative proteomic approach (Dhamad et al., 2016), but their specific contribution in the receptor action still needs to be investigated. Moreover, HSF1 is involved in the regulation of a plethora of non-HSP genes, which support oncogenic processes: cell-cycle regulation, signaling, metabolism, adhesion, and translation (Mendillo et al., 2012). A high level of HSF1 expression was found in cancer cell lines and many human tumors (Vydra et al., 2014) (De Thonel et al., 2011) and was shown to be associated with increased mortality of ER-positive breast cancer patients (Santagata et al., 2011) (Gökmen-Polar and Badve, 2016).

E2-activated HSF1 is transcriptionally potent and takes part in the regulation of several genes essential for breast cancer cell growth and/or ERα action (Vydra et al., 2019). Thus, a hypothetical positive feedback loop between E2/ERα and HSF1 signaling may exist, which putatively supports the growth of estrogen-dependent tumors. Here, to study the cooperation of HSF1 and ESR1 in estrogen signaling and the influence of HSF1 on E2-stimulated transcription and cell growth, we created novel experimental models based on HSF1-deficient cells and performed an in-depth bioinformatics analysis of the relevant genomics data.

## Results

### HSF1 deficiency slows the estrogen-stimulated growth of ERα-positive MCF7 cells

To study the contribution of HSF1 in E2 signaling, we established MCF7 cell lines with reduced HSF1 expression. Firstly, we tested a few *HSF1-*targeting shRNAs (Fig. S1A). Then, the most potent variant that reduced HSF1 level about 10-fold (termed afterward shHSF1) was chosen for further studies. Although the heat shock response was significantly reduced, the expression of *HSP* genes (*HSPA1A*, *HSPH1*, *HSPB1*, and *HSPB8*) was still induced after this HSF1 knockdown (Fig. S1B). Thus, we additionally created MCF7 variants with HSF1 functional knockout using the CRISPR/Cas9 gene targeting approach (termed KO#1 and KO#2 afterward). The complete elimination of HSF1 (Fig. 1A) was connected with a substantial loss of inducibility of *HSP* genes following hyperthermia (Fig. S1B). HSF1 knockdown did not affect the proliferation rate, while the functional HSF1 knockout led to a slight reduction in the proliferation rate under standard conditions (this effect was not visible under less favorable growing conditions, i.e. in 5% dextran-activated charcoal-stripped FBS; Fig. S1C). Also, the increased contribution of cells in the G1 phase was associated with the HSF1 knockout (Fig. S1D) while the ability of cells to form colonies in the clonogenic assay was reduced in both MCF7 experimental models of HSF1 depletion (using shRNA and sgRNA; Fig. 1B). Moreover, the population size of ALDH- positive (stem/progenitor) cells correlated with the HSF1 level and was reduced in HSF1-deficient cells (Fig. 1C). To check if HSF1 deficiency would affect the growth of another ERα-positive cell line, we modified T47D cells using the CRISPR/Cas9 method (Fig. S1E). Under standard conditions, we did not observe differences between unstimulated HSF1+ and HSF1− T47D cell variants in the proliferation and clonogenic assay (not shown). Unlike MCF7 cells, HSF1− T47D cells grew slightly faster than HSF1+ cells but this difference was not statistically significant (Fig. S1F).

**Fig. 1.**
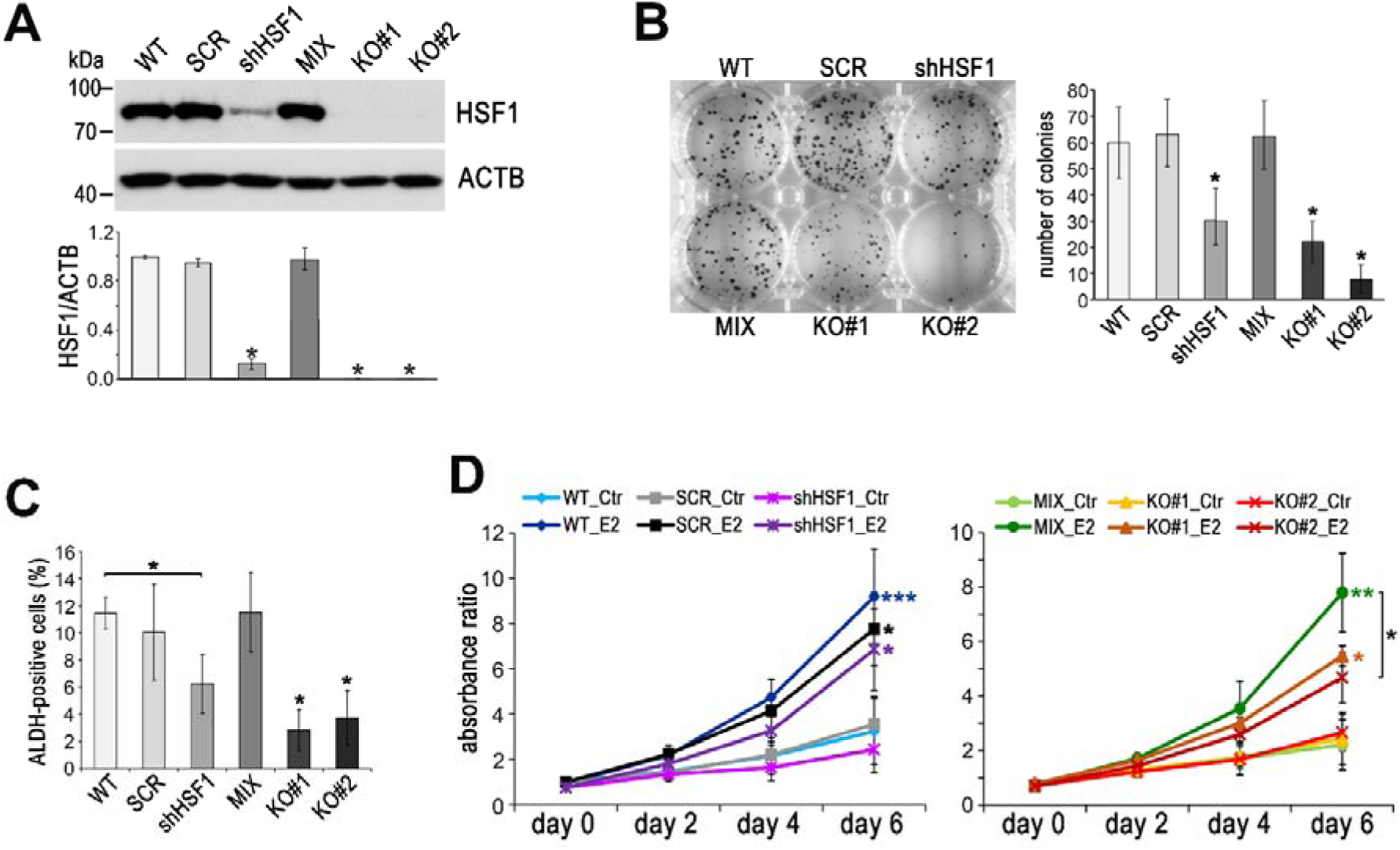
Effect of HSF1 depletion on MCF7 cell growth. **(A)** Western blot analysis of HSF1 level in cell variants: unmodified (WT), stably transduced with non-specific shRNA (SCR), stably transduced with HSF1-specific shRNA (shHSF1), a combination of control clones arisen from single cells following CRISPR/Cas9 gene targeting (MIX), two HSF1 negative clones obtained by CRISPR/Cas9 gene targeting (KO#1, KO#2). Actin (ACTB) was used as a protein loading control. The graph below shows the results of densitometric analysis of HSF1 immunodetection (n=3). *p < 0.05. (**B**) The number of colonies formed by unstimulated cell variants in the clonogenic assay: representative images of single-cell clones stained with crystal violet and their quantification (mean±SD, n=5). *p < 0.05. (**C**) Aldefluor assay of progenitor (ALDH-positive) cell variants assessed by flow cytometry (n=4). *p < 0.05. **(D)** Growth curves of untreated (Ctr) and E2-stimulated cell variants in phenol red-free media with 5% charcoal-stripped FBS assessed using crystal violet staining. Mean and standard deviation from four independent experiments (each in six technical replicates) are shown. ***p < 0.0001, **p < 0.001, *p < 0.05 (next to the curve – compared to the corresponding control, between curves – between cell variants).

We have previously demonstrated that HSF1 was activated after E2 treatment of ERα-positive cells and it was able to bind to the regulatory sequences of several target genes, which correlated with the upregulation of their transcription (Vydra et al., 2019). Since most of these genes code for proteins involved in E2 signaling, we expected that HSF1 downregulation could affect E2-dependent processes, especially cell proliferation. Therefore, we compared E2-stimulated proliferation of HSF1-proficient (WT, SCR, MIX) and HSF1-deficient (shHSF1, KO#1, KO#2) MCF7 cells. The E2-stimulated growth was weaker in the HSF1 knockout cells than in the corresponding control cells but a statistically significant difference was only observed between stimulated KO#2 and MIX cells (Fig. 1D). A similar trend was observed in HSF1 knockdown cells (Fig. 1D). However, E2-stimulated proliferation was not significantly reduced in HSF1 knockout T47D cells (Fig. S1G). These results indicate that HSF1 may influence the growth of ER-positive breast cancer cells, also stimulated by estrogen, although the effect also depends on other factors (differences between cells, culture conditions).

### Transcriptional response to estrogen is inhibited in HSF1-deficient cells

In a search for the mechanism responsible for a distinct response to estrogen in ER-positive cells with different levels of HSF1, we analyzed global gene expression profiles by RNA-seq in MCF7 cell variants (Supplementary Dataset 1). At control conditions (no E2 stimulation), we found relatively few genes differentially expressed in HSF1-proficient (WT, SCR, and MIX) and HSF1-deficient (shHSF1, KO#1, and KO#2) cells that were common for different modes of HSF1 downregulation. These included mainly known HSF1 targets (e.g. *HSPH1*, *HSPE1*, *HSPD1*, *HSP90AA1*) slightly repressed in HSF1-deficient cells. After E2 stimulation, there were 50 genes similarly regulated (47 upregulated and 3 down-regulated) in all HSF1- proficient MCF7 cell variants (Fig. 2A, B). On the other hand, only 13 genes were similarly upregulated after E2 stimulation in all HSF1-deficient MCF7 cell variants (Fig. 2A, C). The geneset enrichment analyses indicated that HSF1 deficiency negatively affected the processes activated by estrogen, especially the early estrogen response (Fig. 2D; terms from other Molecular Signatures Database collections are shown in Fig. S2). Moreover, though almost all genes upregulated by E2 in HSF1-proficient cells were also upregulated in HSF1-deficient cells (except *NAPRT*), the degree of their activation (measured as a fold change E2 vs Ctr) was usually weaker in the latter cells (Fig. 2E), which indicated that the transcriptional response to estrogen was inhibited in the lack of HSF1. Interestingly, however, several E2-dependent genes revealed slightly higher basal expression (without E2 stimulation) in HSF1-deficient cells (Fig. 2F), which suggested that in the absence of E2, HSF1 could be involved in the suppression of these genes.

**Fig. 2.**
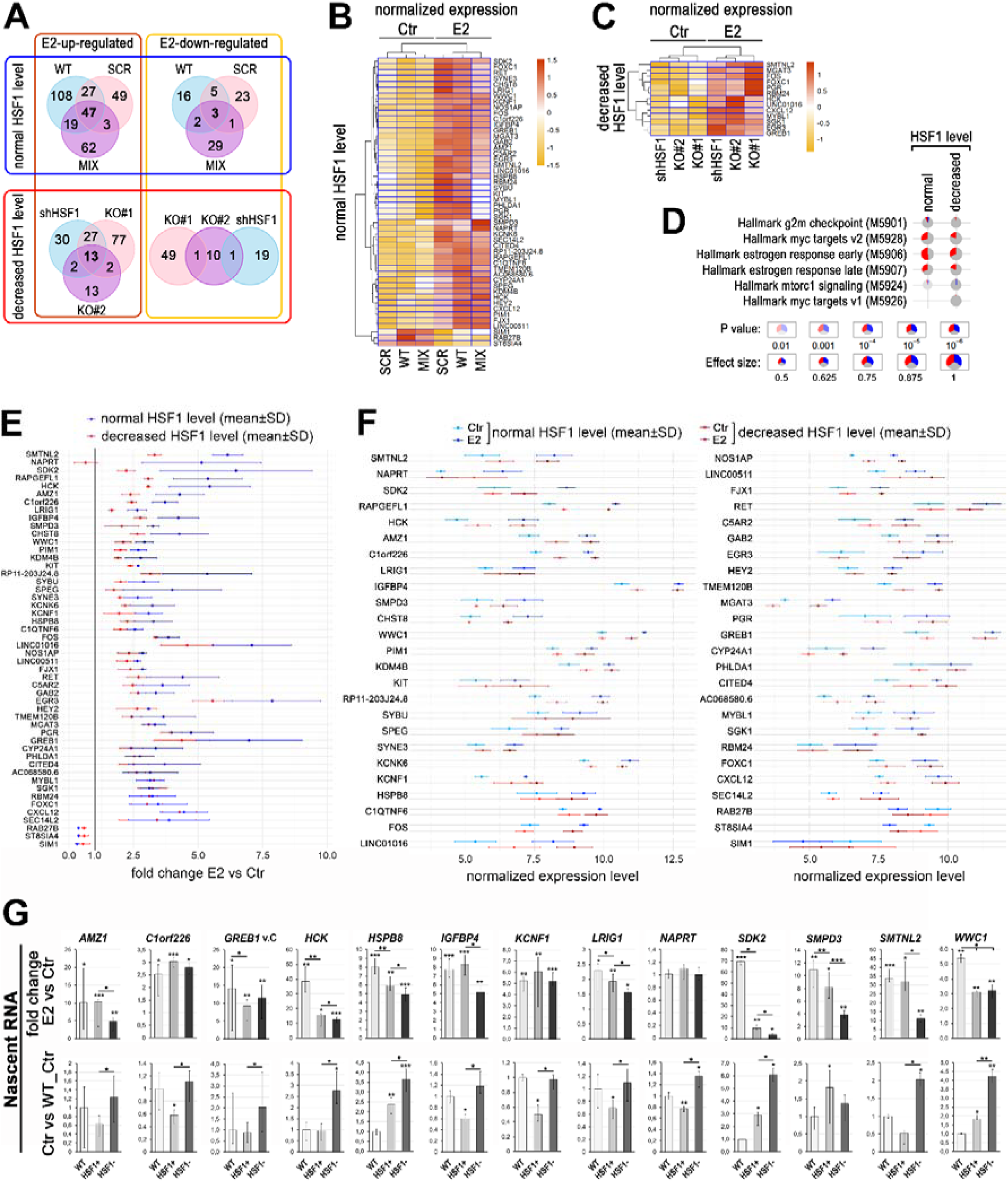
The deficiency of HSF1 reduces a transcriptional response to estrogen (E2) in ER-positive MCF7 cells. (**A**) Overlap of genes and (**B, C**) heatmaps with hierarchical clustering of normalized read counts from RNA-seq (row z-score) for genes stimulated or repressed after the E2 treatment in cells with different levels of HSF1: unmodified (WT), stably transduced with non-specific shRNA (SCR), stably transduced with HSF1- specific shRNA (shHSF1), the combination of control clones arisen from single cells following CRISPR/Cas9 gene targeting (MIX), HSF1 negative single clones obtained by CRISPR/Cas9 gene targeting (KO#1, KO#2). Ctr, untreated cells; E2, 17β-estradiol treatment (10 nM, 4 h). (**D**) Geneset enrichment analysis showing differences between HSF1-proficient and HSF1-deficient cells in response to E2 stimulation. Only significant terms from the hallmark gene sets collection are shown. Blue – a fraction of down-regulated genes, red – fraction of up-regulated genes. (**E**) Comparison of the response to E2 stimulation (E2 vs Ctr) in HSF1-proficient and HSF1-deficient cells. Genes shown in panel B are sorted from the highest to the lowest difference between average fold changes in both cell variants decreased by the standard deviation (SD). Up-regulation – fold change > 1.0, down-regulation – fold change < 1.0. (**F**) Comparison of the expression level (normalized RNA-seq read counts; mean ±SD) of the same set of genes in unstimulated (Ctr) and E2-stimulated HSF1-proficient and HSF1- deficient cells. (**G**) Nascent RNA gene expression analyses by RT-qPCR in MCF7 cells created for validation using DNA-free CRISPR/Cas9 system (HSF1+, six clones with the normal HSF1 level; HSF1−, six HSF1- negative clones). The upper panel shows E2-stimulated changes (E2 vs Ctr fold change; E2 treatment: 10 nM, 4 h), lower panel shows basal expression level represented as fold differences between untreated (Ctr) wild-type, HSF1+, and HSF1− cells. Total RNA analyzes are shown in Fig. S4C. ***p < 0.0001, **p < 0.001, *p < 0.05 (above the bar – compared to the corresponding control, between the bars – between cell variants).

Considering differences between KO#1 and KO#2 HSF1 knockout clones derived from individual cells (similarly, MIX was different from WT cells; Fig. S3), we created an additional experimental model to validate the results described above. Six new individual HSF1-negative (HSF1−) and six HSF1-positive (HSF1+) MCF7 clones obtained using the DNA-free CRISPR/Cas9 system were pooled, characterized for the heat shock response (Fig. S4A), and used for validation analyses. Proliferation tests confirmed that both untreated and E2-stimulated HSF1− cells grew slower than corresponding HSF1+ cells, but the differences were statistically significant only under superior growing conditions (i.e. 10% FBS; Fig. S4B). Out of 13 genes selected for RT-qPCR-based validation using total or nascent RNA, all but *NAPRT* were estrogen-induced (Fig. 2G; Fig. S4C). In the case of 9 (total RNA) or 8 (nascent RNA) genes, the degree of activation was substantially lower in HSF1− than in HSF1+ cells. When the basal expression in E2-untreated cells was compared using the total RNA, 6 genes were expressed at a significantly higher and 1 at a lower level in HSF1− than HSF1+ cells (Fig. S4C). On the other hand, if the nascent RNA was analyzed, there were 12 genes expressed at a higher level in untreated HSF1− cells in comparison to HSF1+ cells (Fig. 2G). Hence, RT-qPCR-based validation analyses generally confirmed differences between HSF1-proficient and HSF1-deficient MCF7 cells revealed by the RNA-seq analyzes.

### HSF1 influences the binding of ESR1 to chromatin

To further study the influence of HSF1 on estrogen signaling, we analyzed ESR1 binding to chromatin in HSF1-proficient and HSF1-deficient MCF7 cells. A list of all ESR1 binding sites detected by ChIP-seq in unstimulated cells and after 30 or 60 minutes of E2 treatment is presented in Supplementary Dataset 2. These analyses revealed that in unstimulated cells, ESR1 binding was more efficient (more binding sites and increased number of tags per peak) in HSF1-deficient cell variant (KO#2) than in corresponding HSF1-proficient control (MIX cells) (Fig. 3A, B) (it is worth noting that the MIX cell variant was also different from wild type cells, indicating that the genome organization was affected by the CRISPR/Cas9 procedure itself, possibly due to off-targets). ESR1 target sequences in *IGFBP4* or *GREB1* are examples of such increased binding efficiency in unstimulated HSF1-deficient cells (Fig. 3D). Estrogen treatment for 30 or 60 minutes resulted in enhanced ESR1 binding in all cell variants. However, fold enrichment (E2 versus Ctr) was lower in HSF1-deficient cells than in HSF1-proficient cells (Fig 3C). Moreover, the number of detected peaks in the E2-treated HSF1-deficient cells was only slightly higher than in unstimulated cells (Fig. 3A) and enhanced ESR1 binding was primarily manifested in sites already existing in unstimulated cells (Fig. 3C, D). We additionally searched for ESR1 binding preferences in HSF1-proficient and HSF1-deficient cells. After estrogen treatment, ESR2 and ESR1 motifs were centrally enriched in ESR1 binding regions in all cell variants (Fig. S5). Moreover, in untreated cells, the motif for PBX1 (not centrally enriched in peak regions), which is a pioneer factor known to bind to the chromatin before ESR1 recruitment (Magnani et al., 2011), was identified by MEME-ChIP analysis in all cell variants (not shown). This indicates that ESR1 chromatin binding preferences were not substantially changed in HSF1-deficient cells.

**Fig. 3.**
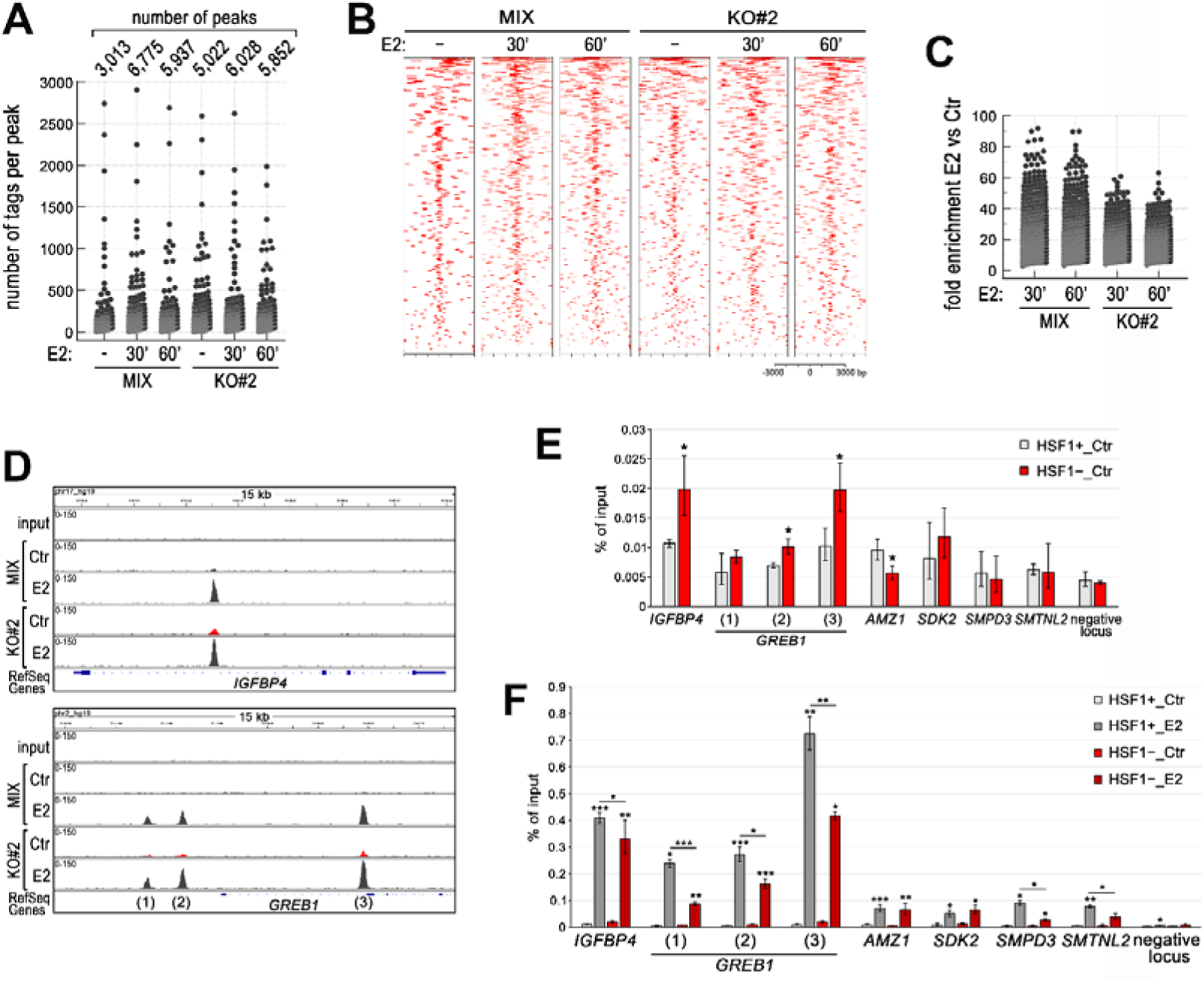
HSF1 deficiency influences the binding of ESR1 to chromatin in ER-positive MCF7 cells. (**A**) Number of peaks and peak size distribution (number of tags per peak), (**B**) heatmap visualization of ESR1 ChIP-seq data (versus input), and (**C**) binding enrichment (fold enrichment E2 versus Ctr) after E2 stimulation (10 nM for 30 or 60 minutes) in HSF1-deficient cells (KO#2) and corresponding control (MIX, a combination of control clones arisen from single cells following CRISPR/Cas9 gene targeting). Heatmaps depict all ESR1 binding events centered on the peak region within a 3 kb window around the peak. Peaks in each sample were ranked on intensity. (**D**) ESR1 peaks identified by MACS in ChIP-seq analyses and visualized by the IGV browser in unstimulated cells (Ctr) and after E2 treatment (10 nM, 30 min). The scale for each sample is shown in the left corner. Peaks showing more efficient local ESR1 binding in untreated HSF1-deficient cells are marked in red. (**E**) Comparison of ESR1 binding efficiency (by ChIP-qPCR) in selected sequences in untreated MCF7 cells created for validation using DNA-free CRISPR/Cas9 system: HSF1+, six clones with the normal HSF1 level; HSF1−, six HSF1-negative clones. (**F**) ESR1 binding after E2 stimulation: Ctr, untreated cells; E2, 17β-estradiol treatment (10 nM, 30 min). ***p < 0.0001, **p < 0.001, *p < 0.05 (above the bar – compared to the corresponding control, between the bars – between cell variants).

To validate ChIP-seq results, we analyzed the influence of HSF1 on the binding of ESR1 to selected target sites by ChIP-qPCR using the novel MCF7 CRISPR/Cas9 model. In the case of *IGFBP4* and *GREB1* (i.e. sequences highly enriched with ESR1 after E2 stimulation), the binding efficiency of ESR1 (shown as a percent of input) was higher in unstimulated HSF1− cells than in corresponding HSF1+ cells (Fig. 3E). On the other hand, although estrogen treatment strongly induced ESR1 binding, this induction was considerably lower in HSF1− cells (Fig. 3F). Therefore, we validated ChIP-seq results and confirmed that in strongly-responsive ESR1 binding sites deficiency of HSF1 correlated with enhanced binding of unliganded ESR1 and weaker enrichment of ESR1 binding upon estrogen stimulation. However, other patterns of the response are also possible, especially in sequences that were weakly enriched in ESR1 after stimulation, as exemplified by *AMZ1*, *SDK2*, *SMPD3*, and *SMTNL2* (Fig. 3E, F).

### HSF1 deficiency is associated with altered interactions between HSP90 and ESR1

ESR1 is known to be kept in an inactive state by HSP90 (Pratt and Toft, 1997), in particular by HSP90AA1 (Dhamad et al., 2016) that is the HSF1 transcriptional target. Thus, looking for a reason for the dysregulated ESR1 binding in HSF1-deficient cells we focused on ESR1 and HSP90 interactions. Analyzes of the proximity of both proteins by PLA revealed that the number of ESR1/HSP90 complexes decreased after estrogen treatment in HSF1+ MCF7 cells (Fig. 4A). This indicates that liganded (and transcriptionally active) ESR1 is indeed released from the inhibitory complex with HSP90. *HSP90AA1* expression was substantially reduced in HSF1-deficient cells (RNA-seq analyses), which correlated with the reduced HSP90 protein level. Also, the ESR1 level was considerably decreased in most HSF1-deficient cell variants (except KO # 1 cells; not shown), especially in cells cultured in phenol-free media (Fig. 4B, Fig. S4A). Therefore, we hypothesized that the number of ESR1/HSP90 complexes could be reduced in HSF1-deficient cells, which would result in enhanced basal transcriptional activity of ESR1 in untreated cells. However, we observed an increased number of such complexes both in untreated and E2-stimulated HSF1-deficient cells when compared to HSF1-proficient cells (Fig. 4A). This indicates that the response to estrogen could be dysregulated in HSF1-deficient cells, also at the level of ESR1/HSP90 interactions, in a mechanism not related directly to the HSP90 and ESR1 downregulation mediated by the HSF1 deficiency.

**Fig. 4.**
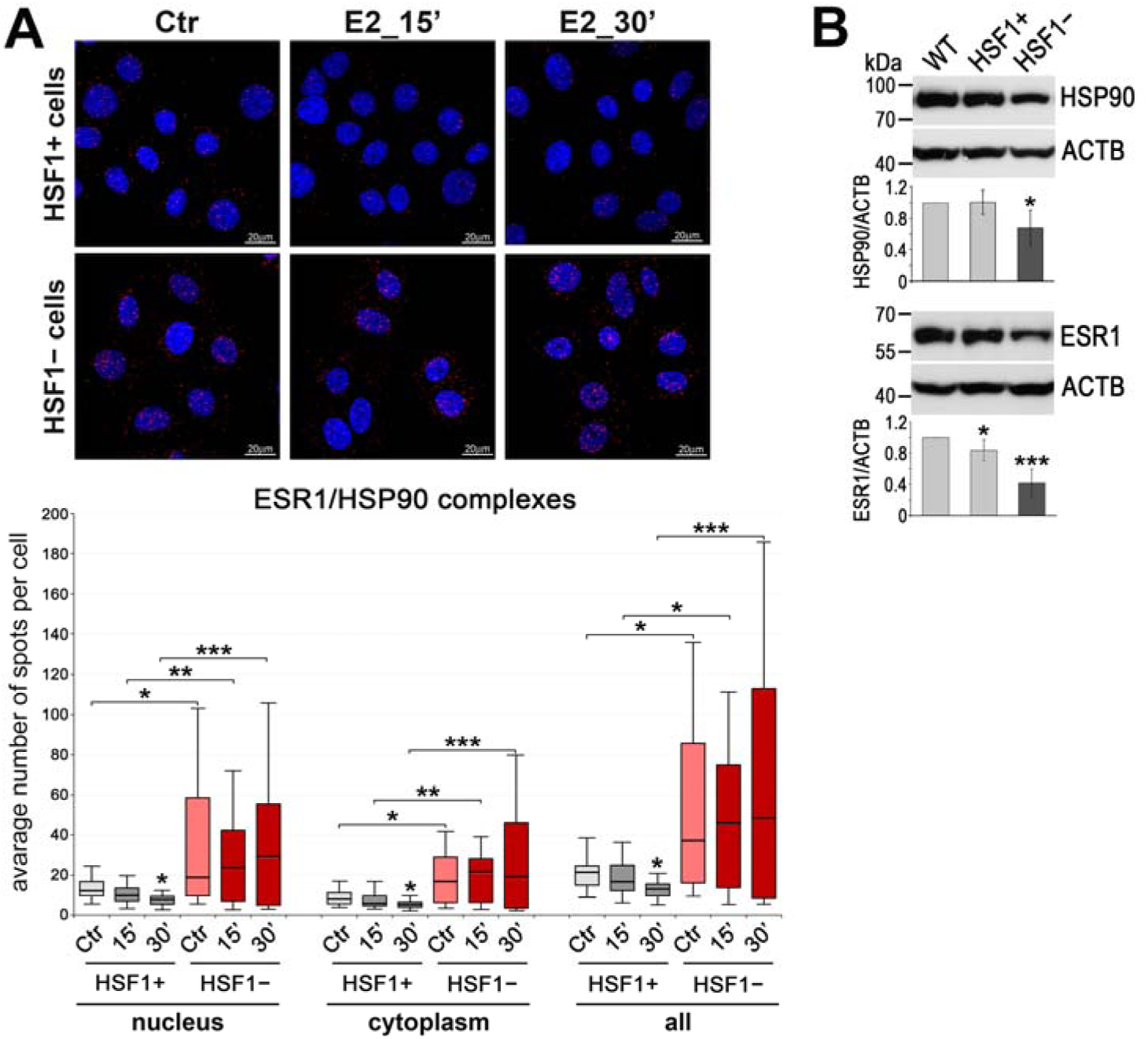
HSF1-deficiency is associated with the reduced HSP90 and ESR1 levels and altered HSP90/ESR1 interactions. (**A**) Interactions between ESR1 and HSP90 assessed by Proximity Ligation Assay (PLA; red spots) in HSF1-positive (HSF1+) and HSF1-negative (HSF1−) MCF7 cells created using DNA-free CRISPR/Cas9 system. Ctr, untreated cells; E2, 17β-estradiol treatment (10 nM). DNA was stained with DAPI. Scale bar, 20 μm. The mean number of spots per cell is shown in the boxplot below. The line dividing the box represents the median and the upper and lower side of the box shows the upper and lower quartiles, respectively. The whiskers show the highest and lowest values. ***p < 0.0001, **p < 0.001, *p < 0.05 (above the bar – compared to the corresponding control, between the bars – between cell variants). For detection of HSP90 and ESR1 by immunofluorescence see Fig. S6. (**B**) HSP90 and ESR1 levels assessed by western blot. Actin (ACTB) was used as a protein loading control. The graphs show the results of densitometric analyses (n=4). ***p < 0.0001, *p < 0.05.

### HSF1 can cooperate with ESR1 in chromatin binding and participate in the spatial organization of chromatin loops

Since estrogen-activated HSF1 has been shown to bind to chromatin, we compared the binding patterns of ESR1 and HSF1 in wild-type MCF7 cells (using our ChIP-seq data deposited in the NCBI GEO database; acc. no. GSE137558; (Vydra et al., 2019)). Although in untreated cells (Ctr) there were 1,535 and 2,248 annotated peaks for ESR1 and HSF1 respectively (compared to the input), only a few (below 50) binding sites with overlapped peaks for both transcription factors were identified. Moreover, these common binding regions were characterized by a small number of tags (smaller in the case of ESR1) (Fig. 5A; Supplementary Dataset 3, sheet 1). On the other hand, the search for ESR1 and HSF1 common binding regions created after estrogen treatment (E2 vs Ctr) returned more than 200 peaks (Supplementary Dataset 3, sheet 2). Numbers of tags per peak and fold enrichment increased after E2 stimulation for both factors yet more for ESR1 than HSF1 binding in such regions (Fig. 5B). Although distributions of the fold enrichment after E2 stimulation in all peaks for each transcription factor separately and the overlapped ones were similar, sites mapped to the common binding regions represented only a small fraction of the total number of ESR1 binding sites (∼2.5% from 8,320 peaks; in the case of HSF1 this represents 35% of 571 peaks) (Fig. 5C). These results suggest that co-binding of both factors in the same DNA region is not critical in the regulation of the ESR1 transcriptional activity. Instead, we postulate that HSF1 may influence the organization of the chromatin loops created after estrogen stimulation. There were different patterns of estrogen-stimulated binding of ESR1 and HSF1 to chromatin (Fig. S7). Generally, more abundant ESR1 binding was observed at overlapped sites (e.g. in the case of *GREB1*; Fig. 5D and Fig. S7A), but HSF1 binding could be stronger as well (Fig. S7B). When we combined ESR1 and HSF1 ChIP-seq peaks with data from chromatin interaction analysis by paired-end tag sequencing (ChIA-PET) performed by (Fullwood et al., 2009), it was evident that the HSF1-binding sites mapped to ESR1-interacting loci (ESR1 anchor regions) even if actual ESR1 binding was not detected in the same locus (examples of such anchors in *FAM102A*, *HSPB8*, *PRKCE*, and *WWC1* regulatory sequences are shown in Fig 5D). HSF1 peaks unrelated to ESR1 anchoring were also existing (Fig. S7C). Further analyzes of the spatial organization of chromatin by chromosome conformation capture (3C) technique revealed that some interactions between different ESR1 anchor regions were dependent on the presence of HSF1. This is exemplified by *HSPB8* and *WWC1* loci analyzed in HSF1-proficient and HSF1- deficient cells (Fig. 5E), which confirms the role of HSF1 in the formation of ESR1-mediated chromatin loops.

**Fig. 5.**
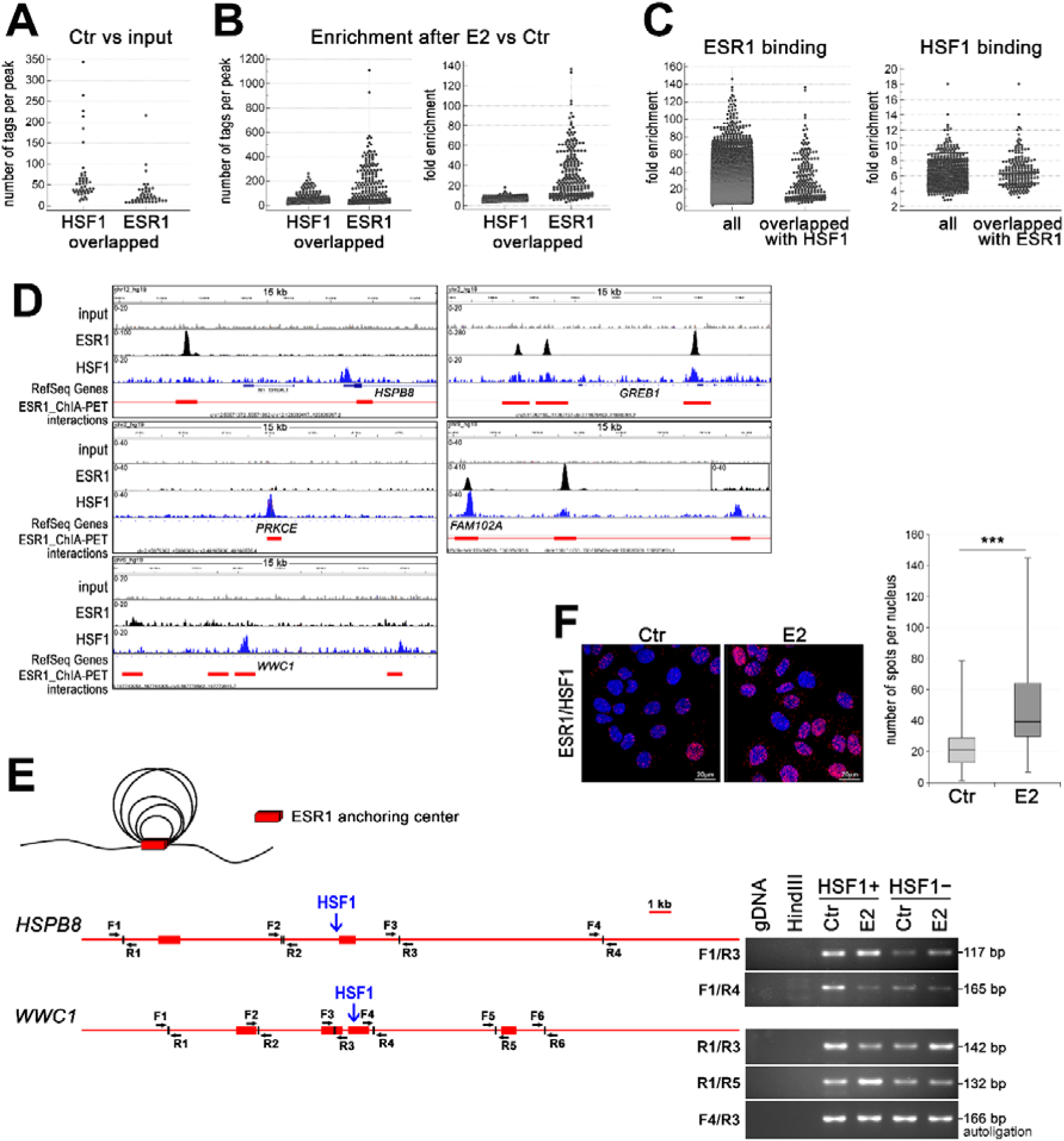
HSF1 may cooperate with ESR1 in chromatin binding and take a part in chromatin organization. (**A**) Overlapped HSF1 and ESR1 ChIP-seq peaks in untreated wild-type MCF7 analyzed for peak size distribution (number of tags per peak). (**B**) Overlapped HSF1 and ESR1 ChIP-seq peaks in wild-type MCF7 after E2 stimulation analyzed for peak size distribution (number of tags per peak) and fold enrichment. (**C**) Comparison of the binding enrichment (E2 vs Ctr) of all ESR1 and HSF1 peaks and overlapped peaks identified after stimulation in wild-type MCF7 cells. (**D**) Examples of ESR1 and HSF1 peaks identified by MACS in ChIP-seq analyses in wild-type MCF7 cells after E2 treatment and corresponding ChIA-PET interactions (Fullwood et al., 2009) downloaded from ENCODE database and visualized by the IGV browser. The red bar shows the ESR1 anchor region (interacting loci), red line – the intermediate genomic span between the two anchors forming a putative loop; the scale for each sample is shown in the left corner. (**E**) ESR1-mediated chromatin interactions analyzed by chromosome conformation capture (3C) technique in *HSPB8* and *WWC1* loci. The scheme represents ESR1 anchor regions (red bars), HSF1 binding sites (blue arrows), and forward (F) and reverse (R) primers around subsequent *HindIII* cleavage sites. A model of chromatin loops resulting from interactions between ESR1 anchor regions is also illustrated above. Interactions between selected DNA regions were analyzed by PCR in untreated and E2-stimulated HSF1+ and HSF1− cells. (**F**) Interactions between ESR1 and HSF1 assessed by PLA (red spots) in wild-type MCF7 cells after E2 treatment. DNA was stained with DAPI. Scale bar, 20 μm. The number of spots per nucleus is shown (boxplots represent the median, upper and lower quartiles, maximum and minimum. ***p < 0.0001. E2, 10 nM for 60 minutes (or 30 minutes for 3C).

Though the co-binding of HSF1 and ESR1 to DNA was rare and relatively weak, particularly in untreated cells, the proximity of both factors was easily detected. In general, both transcription factors co-localized in the nucleus when assessed by immunofluorescence (Fig. S6). Thus, PLA spots indicating putative HSF1/ESR1 interactions were mainly located in the nucleus and their number dramatically increased after E2 treatment (Fig. 5F). However, large diversity was observed between individual cells, which suggests that also HSF1 binding to DNA may be differentiated at the single-cell level. Nevertheless, we concluded that the proximity of HSF1 and ESR1 putatively reflecting their interactions frequently happens in the cell nucleus.

### The combined expression level of ESR1 and HSF1 can be used to predict the survival of breast cancer patients

Our *in vitro* analyses indicated that HSF1 could support the transcriptional action of ESR1 upon estrogen treatment. On the other hand, HSF1-regulated chaperones are necessary to keep estrogen receptors in an inactive state in the absence of ligands, which collectively indicated important functional crosstalk between both factors. Therefore, to further study the significance of the interaction between ESR1 and HSF1 in actual breast cancer, we utilized RNA-seq data deposited in the TCGA database. The analysis revealed that the transcript level of *HSF1* negatively correlated with the *ESR1* transcript level, although this tendency was relatively weak (Fig. 6A). Neither *ESR1* nor *HSF1* transcript levels analyzed separately had a significant prognostic value (Fig. S8). Therefore, out of all breast cancer cases, we selected four groups characterized by different levels of ESR1 (mRNA and protein level) and *HSF1* (mRNA) expression: ER−/HSF1^low^, ER−/HSF1^high^, ER+/HSF1^low^, and ER+/HSF1^high^ (Fig. 6B). These groups varied in molecular subtypes composition. In ER+ cancers (luminal A, luminal B, and normal-like), the HSF1^low^ group was more homogenous (mostly luminal A) than the HSF1^high^ group. In ER– cases (basal-like and HER2-enriched), the HSF1^high^ group was more homogenous (mostly basal-like) (Fig. 6C). Analyses of the survival probability showed that a high *HSF1* expression had a greater negative effect on the survival of ER– than ER+ cancers. The most divergent groups were: ER+/HSF1^low^ and ER−/HSF1^high^ (better and worse prognosis, respectively; p=0.0044), which represented luminal A and basal-like enriched groups (Fig. 6D). These analyses indicate that *HSF1* and *ESR1* may have an additive effect on survival and have prognostic value only if analyzed together. The difference between ER+/HSF1^low^ and ER−/HSF1^high^ cancers was also clearly visible in the multidimensional scaling (MDS) plots where the cancer cases belonging to these groups were separated. MDS plotting generally separated ER+ cases from ER−/HSF1^high^ cases, while ER−/HSF1^low^ cases were scattered between them (Fig. 6E). On the other hand, HSF1^high^ and HSF1^low^ cases were not separated although they were slightly shifted against each other. When looking at molecular subtypes, it became apparent that ER−/HER2-positive cancers were separated from ER−/basal-like cancers and slightly overlapped with ER+ cancers.

**Fig. 6.**
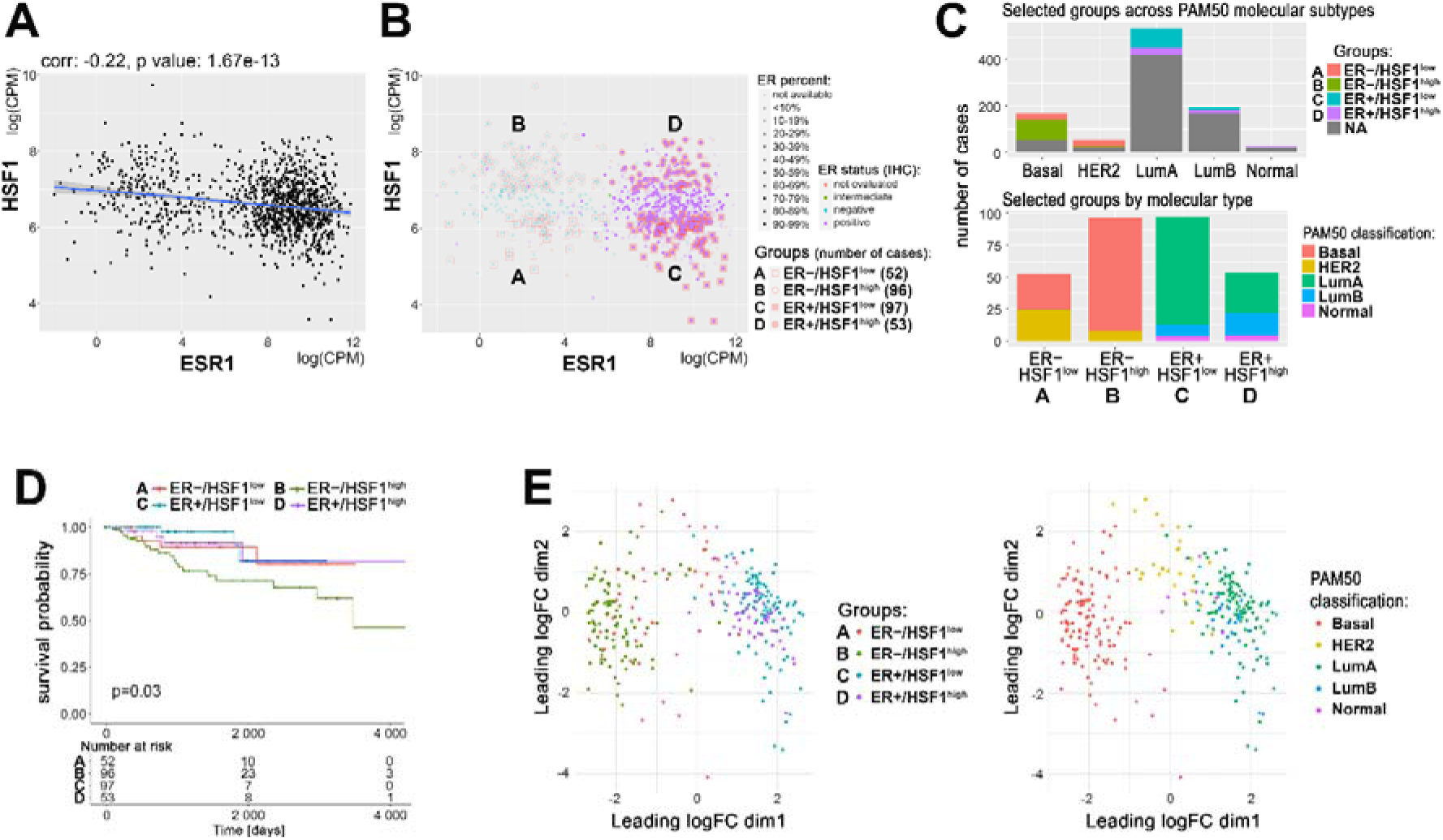
Relationship between ESR1 and HSF1 expression in breast cancer. (**A**) Correlation of *HSF1* and *ESR1* transcript level in all TCGA breast cancers. Each spot represents one cancer case; log(CPM), log2-counts per million. (**B**) Groups with different mRNA levels of *HSF1* and *ESR1* selected for further analyses. In the case of ESR1, also the protein level assessed by immunohistochemistry (IHC) was taken into consideration. (**C**) Characteristics of selected groups in relation to the molecular subtypes of breast cancer. (**D**) Kaplan-Meier plots for all selected groups. (**E**) MDS plots of selected cases with marked: ER and HSF1 statuses (left) or molecular subtypes (right). ER+/−, estrogen receptor-positive/negative; HSF1^high^, high *HSF1* level, HSF1^low^, low *HSF1* level.

### HSF1 increases the diversity of the transcriptome in ER-positive breast cancers

Furthermore, we analyzed global gene expression profiles in breast cancers with different ESR1 and HSF1 statuses. Differential expression tests between the above-selected groups of patients (Supplementary Dataset 4) revealed that generally, ESR1 had a much stronger influence on the transcriptome (i.e., ER+ versus ER−) than HSF1 (i.e., HSF1^high^ versus HSF1^low^). Nevertheless, differences between ER+ and ER− cases were higher in the presence of high levels of HSF1, which implicates that HSF1 increases the diversity of the transcriptome of ER+ cancers. Also, the differences in the transcript levels between HSF1^high^ and HSF1^low^ cancers were higher in ER+ than ER− cases (Fig. 7A). Remarkably, the most divergent were ER+/HSF1^low^ and ER−/HSF1^high^ cancers, which resembled the most significant differences in the survival probability (Fig. 6D). Then, we looked at differences in numbers of differently expressed genes (DEGs) between patients’ groups. To eliminate the possible influence of the group size on DEGs, we repeated each test 10 times, randomly subsampling groups to an equal number of cases and averaging the number of DEGs. Furthermore, to check whether heterogeneity of selected groups regarding molecular subtypes could affect observed differences in gene expression profiles, only basal-like (ER−) and luminal A (ER+) cancers were included in these tests (Fig. 7B). In general, these analyses also revealed that the number of genes differentiating ER+ and ER− cases were higher in HSF1^high^ cancers, while the number of genes differentiating HSF1^high^ and HSF1^low^ cases was higher in ER+ cancers. The most divergent were again ER+/HSF1^low^ and ER−/HSF1^high^ cases while the most similar, ER−/HSF1^low^ and ER−/HSF1^high^ (Fig. 7C). This tendency was maintained when groups with mixed molecular subtypes composition were analyzed as well as more homogenous cancer groups (i.e., only basal-like and luminal A). Differences in gene expression profiles between pairwise compared groups of cancer are further illustrated on volcano plots that additionally separated upregulated and downregulated genes (Fig. S9). We further searched for the hypothetical influence of the HSF1 status on functions of ESR1-related genes in actual cancer tissue. The geneset enrichment analysis identified terms related to estrogen response among the most significant ones associated with transcripts differentiating between ER+ and ER− cancers (Fig. S10). The more detailed analysis focused on terms related to hormone signaling and metabolism showed differences between HSF1^high^ and HSF1^low^ cases when ER+ and ER− cancers were compared. These analyses indicate that HSF1 may enhance estrogen signaling. On the other hand, the analysis focused on terms related to response to stimulus and protein processing (i.e., functions presumed to be dependent on HSF1 action via the HSPs expression), revealed that most of them reached the statistical significance of differences between ER+/HSF1^high^ and ER−/HSF1^high^ cases (Fig. 7D).

**Fig. 7.**
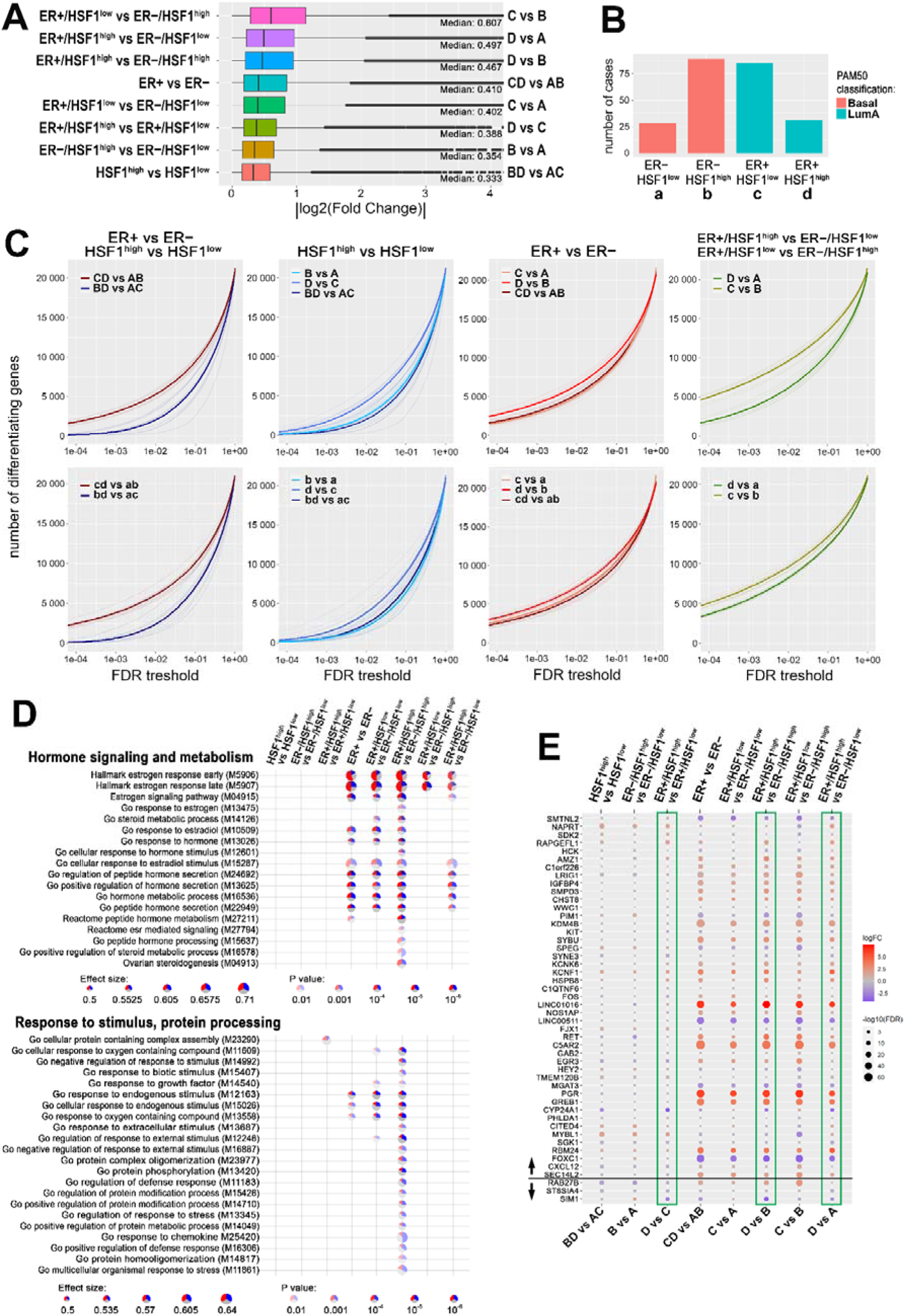
HSF1 increases the diversity of the transcriptome of ER-positive breast cancer. (**A**) Boxplots of fold changes (logFC absolute values) illustrating differences in gene expression between groups characterized in Fig. 6. The line dividing the box represents the median of the data and the right and left side of the box shows the upper and lower quartiles respectively. The whiskers show the highest and lowest values, excluding outliers, which are shown as circles. (**B**) Composition of ER+ and ER− groups with different levels of HSF1 reduced to one molecular subtype. (**C**) The number of differently expressed genes (y-axis) plotted cumulatively against the False Discovery Rate value of differences (x-axis). Comparisons of ER+ and ER− cancer cases as well as HSF1^high^ and HSF1^low^ cases (upper graphs; all cases, for group indexes see Panel A) and cases from pre-selected cancer types (lower graphs; for group indexes see Panel B). (**D**) Geneset enrichment analyses showing differences between ER+ and ER− breast cancers with different HSF1 levels. Terms related to hormone signaling and metabolism and response to stimulus and protein processing in comparisons between groups selected in Fig. 6B. Blue – a fraction of down-regulated genes, red – fraction of up-regulated genes. (**E**) Differences in the expression of the E2-regulated gene set (as identified in MCF7 cells by RNA-seq; see Fig. 2) between breast cancers with different levels of ESR1 and HSF1 selected from the TCGA database and qualified to four groups as shown in Fig. 6B. FDR, False Discovery Rate; logFC, log fold change. Green boxes mark all possible comparisons between the ER+/HSF1^high^ group to other groups. The black horizontal line separates genes up- and down-regulated after E2 treatment in MCF7 cells.

We additionally compared the expression of E2-regulated genes (the set identified in MCF7 cells by RNA-seq, i.e. 47 up-regulated and 3 down-regulated genes; Fig. 2) in selected groups of breast cancers with different levels of ESR1 and HSF1. The analysis revealed the highest up-regulation of *PGR* and *LINC01016* genes in ER+ compared to ER− cancers (regardless of HSF1 status) (Fig. 7E). It is noteworthy, however, that not all genes up-regulated by E2 in MCF7 cells revealed an increased expression level in ER+ compared to ER− cancers. Especially, *FOXC1* and *LINC00511* were expressed at a higher level in ER− cancers. Moreover, regardless of ER status, cancers with high HSF1 levels revealed a higher expression of *MYBL1* and *NAPRT* than cancers with low HSF1 levels. Furthermore, expression of few genes systematically differentiated cancers with high levels of both factors (ER+/HSF1^high^) compared to cancers with the low level of at least one factor (including *RAPGEFL1*, *AMZ1*, *KCNF1*, *HSPB8* up-regulated, and *CYP24A1, SIM1* down-regulated in ER+/HSF1^high^ cancers), which was consistent in all relevant comparisons (marked with green boxes in Fig. 7E). Nevertheless, the observed features of gene expression profiles confirmed collectively that HSF1 affects the genomic action of ESR1 in breast cancer.

## Discussion

The precise mechanisms by which estrogens stimulate the proliferation of breast cancer cells are still unclear. We found that HSF1-deficiency in ER-positive breast cancer cells could slow down the mitogenic effects of estrogen. This may be a consequence of a reduced transcriptional response to estrogen in these cells and therefore implies that HSF1 may support estrogen action. Indeed, analyses of the transcriptome of breast cancers from the TCGA database showed higher transcriptome diversity in ER-positive cases with high expression of *HSF1* than with low *HSF1* levels. High HSF1 nuclear levels (estimated by immunohistochemistry in patients with invasive breast cancer at diagnosis; in situ carcinoma only and stage IV breast cancer were excluded from outcome analysis) were previously associated with decreased survival specifically in ER-positive breast cancer patients (Santagata et al., 2011). However, in another study performed on samples from patients with ER-positive tumors, only a weak association was found between the HSF1 protein expression and poor prognosis (Gökmen-Polar and Badve, 2016). Nevertheless, both studies showed a significant correlation between *HSF1* transcript levels and the survival in ER-positive breast cancer patients. In our analysis, using non-preselected data from the TCGA gene expression database, we found that although high *HSF1* levels slightly reduced the survival in ER-positive cancer patients, they had a greater negative outcome on survival in ER- negative patients. Therefore, we concluded that *HSF1* had prognostic value when analyzed together with *ESR1* transcript level.

The mechanism of supportive HSF1 activity in ER-positive cells was already proposed, by which upon E2 treatment, HSF1 is phosphorylated via ESR1/MAPK signaling, gains transcriptional competence, and activates several genes essential for breast cancer cell growth and/or ESR1 action (Vydra et al., 2019). Here we found that HSF1-deficiency results in a weaker response to estrogen stimulus of many estrogen-induced genes. It is noteworthy, that the reduced transcriptional response to estrogen could at least partially result from the enhanced binding of unliganded ESR1 to chromatin and higher basal expression of ESR1-regulated genes. This suggests that HSF1-dependent mechanisms may amplify ESR1 action upon estrogen stimulation while inhibiting it in the absence of ligands. The proper action of ERs depends on HSF1-regulated chaperones, especially HSP90. As expected, the number of HSP90/ESR1 complexes decreased after ligand (E2) binding in cells with the normal level of HSF1. However, although HSP90 was down-regulated in HSF1-deficient cells, more HSP90/ESR1 complexes were found both in untreated and estrogen-stimulated cells. Hence, increased activity of ESR1 in HSF1-deficient cells could not be explained by reduced sequestration of unliganded ESR1 by HSP90. Accordingly, additional HSF1-dependent factors may influence formation of these complexes. Ligand-independent genomic actions of ESR1 are also regulated by growth factors that activate protein-kinase cascades, leading to phosphorylation and activation of nuclear ERs at EREs (Stellato et al., 2016). Involvement of HSF1 in the repression of estrogen-dependent transcription was reported in MCF7 cells treated with heregulin (NRG1), the ligand for the NEU/ERBB2 receptor tyrosine kinase (Khaleque et al., 2008). Interactions of HSF1 with the co-repressor metastasis-associated protein 1 (MTA1) and several additional chromatin-modulating proteins were implicated in that process. Therefore, it cannot be ruled out that HSF1 influences unliganded and liganded ESR1 by various mechanisms that have to be further investigated.

Transcriptional activation by ESR1 is a multistep process modulated by coactivators and corepressors. Cofactors interact with the receptor in a ligand-dependent manner and are often part of large multiprotein complexes that control transcription by recruiting components of the basal transcription machinery, regulating chromatin structure, and/or modifying histones (Welboren et al., 2009) (Kovács et al., 2020) (Pescatori et al., 2021). Liganded ESR1 may bind directly to DNA (to ERE), and indirectly via tethering to other transcription factors such as FOS/JUN (AP1), STATs, ATF2/JUN, SP1, NFκB (Björnström and Sjöberg, 2005) (Welboren et al., 2009) (Heldring et al., 2011). Here we found that HSF1 can potentially be an additional factor tethering liganded ESR1 to DNA. ESR1 has been shown to function via extensive chromatin looping to bring genes together for coordinated transcriptional regulation (Fullwood et al., 2009). Since ESR1 anchor sites were identified also in sites bound by HSF1 but not ESR1, we propose that HSF1 may be a part of this “looping” machinery (other components in the same anchoring center are also possible). In general, estrogen-induced HSF1 binding was weaker than ESR1 binding. However, PLA analyses indicated a large heterogeneity in a cell population regarding ESR1 and HSF1 interactions and showed individual cells in which such interactions induced by estrogen were very strong. The final transcriptional activity of ESR1 is modulated by interactions with various tethering factors, including HSF1. Therefore we hypothesize that it can be modulated differently at the single cell level by different cofactors and chromatin remodeling factors. Thus, the response measured on the whole cell population is heterogeneous and stochastic when a single cell is considered.

Estrogen-dependent cancers are treated with hormonal therapies, and high levels of HSF1 have been associated with antiestrogen resistance (Silveira et al., 2021). It was proposed that overexpression of HSF1 in ER-positive breast cancers was associated with a decreased dependency on the ERα-controlled transcriptional program for cancer growth. However, this conclusion was based on experiments performed without estrogen stimulation. Our *in vitro* studies indicate that the influence of HSF1 on ESR1 action depends on the presence of the estrogen and HSF1 may repress ERα-controlled transcriptional program only in the absence of the ligand. Enhanced resistance to hormonal therapies could be mediated by HSF1-regulated genes. Heat shock proteins themselves can be prognostic factors in breast cancer and especially oncogenic properties of HSP90AA1 correlated with aggressive clinicopathological features and resistance to the treatment (Whitesell et al., 2014) (Klimczak et al., 2019). Here we proposed a novel mechanism of the HSF1 action in ER-positive breast cancers, which is independent of typical HSF1-regulated genes. This mechanism assumes that HSF1 influences the transcriptional response to estrogen via the re-organization of chromatin structure in estrogen-responsive genes. This mode of HSF1 action may be important in all ESR1-expressing cells. ESR1 is for example a critical transcription factor that regulates epithelial cell proliferation and ductal morphogenesis during postnatal mammary gland development. It is noteworthy that HSF1 has been shown to promote mammary gland morphogenesis by protecting mammary epithelial cells from apoptosis and increasing their proliferative capacity (Xi et al., 2012).

In conclusion, HSF1 and ESR1 cooperate in response to estrogen stimulation. Estrogen via ESR1 and MAPK activates HSF1 (Vydra et al., 2019) which together with ESR1 forms new chromatin loops that enhance estrogen-stimulated transcription (Fig. 8). Moreover, HSF1 may be involved in repression of unliganded ESR1. Genes activated by ESR1 and HSF1 play an important role in regulating the growth of estrogen-dependent tumors and the combination of both factors has a prognostic value in breast cancer patients.

**Fig. 8.**
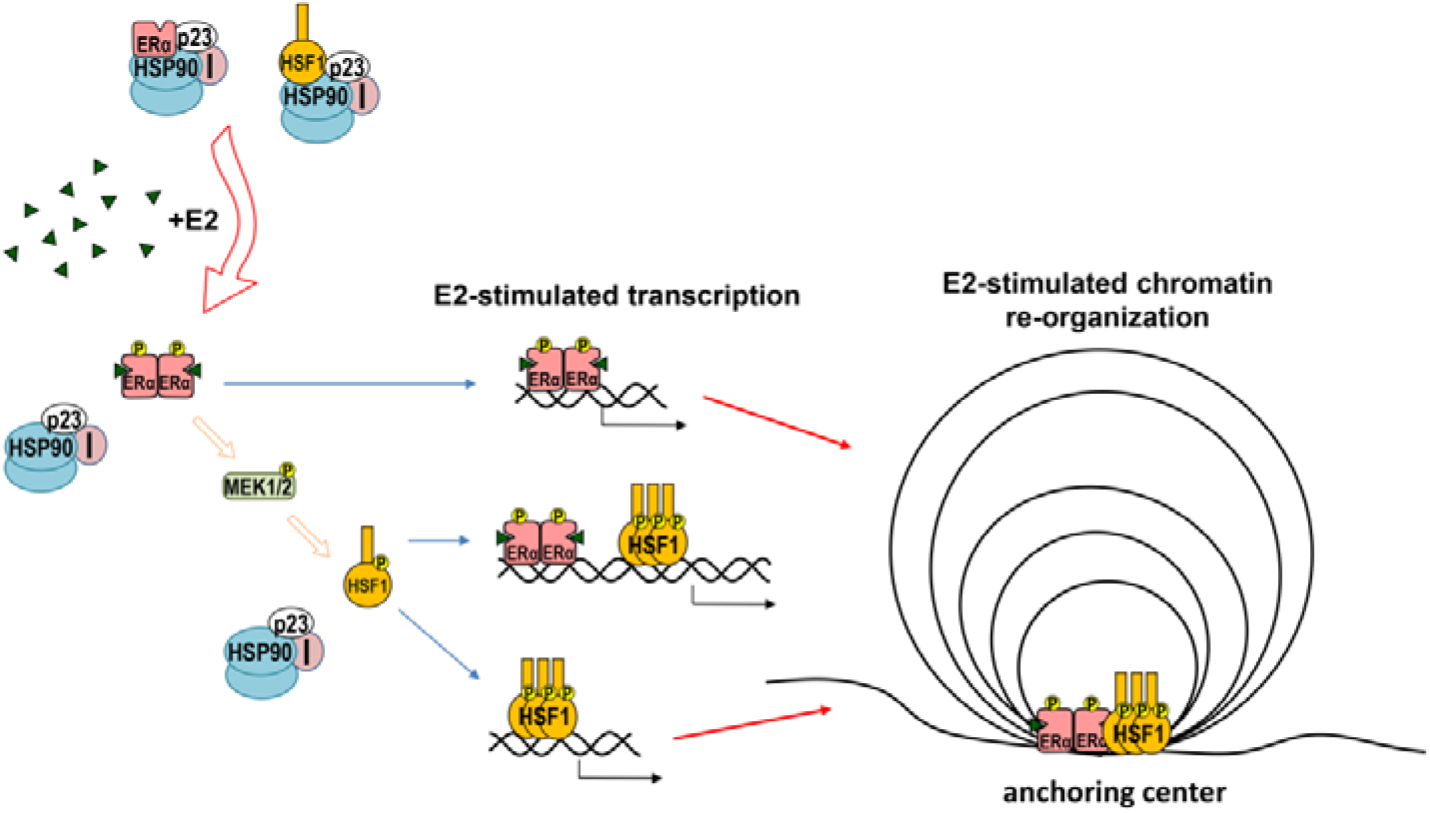
Model of cooperation between ESR1 (ERα) and HSF1 in response to estrogen (E2) stimulation. Both ESR1 and HSF1 are kept in an inactive state by the complexes of HSP90, p23, and immunophilins (I). Binding of E2 to ESR1 is connected with the release of the chaperone complex and activation of ESR1 leading to the phosphorylation of MEK1/2 followed by HSF1 activation. Oligomers of active transcription factors can bind to DNA and cooperate in the regulation of the transcription either directly or through chromatin re-organization.

## Materials and Methods

### Cell lines and treatments

Human MCF7 and T47D ERα-positive breast cancer cell lines were purchased from the American Type Culture Collection (ATCC, Manassas, VA, USA) and the European Collection of Authenticated Cell Cultures (ECACC, Porton Down, UK), respectively. Cells were cultured in DMEM/F12 medium (Merck KGaA, Darmstadt, Germany) supplemented with 10% fetal bovine serum (FBS) (EURx, Gdansk, Poland). For heat shock, logarithmically growing cells were placed in a water bath at a temperature of 43 °C for one hour. The cells were allowed to recover for the indicated time in a CO_2_ incubator at 37 °C. For estrogen treatment, cells were seeded on plates and the next day the medium was replaced into a phenol-free medium supplemented with 5% or 10% dextran-activated charcoal-stripped FBS (PAN-Biotech GmbH, Aidenbach, Germany). 17β-estradiol (E2; Merck KGaA) was added 48 hours later to a final concentration of 10 nM (an equal volume of ethanol was added as vehicle control) for the indicated time. For longer E2 treatments, the medium was changed every two days. The growth media were not replaced either before or after treatments. Cells were routinely tested for mycoplasma contamination.

### HSF1 down-regulation using shRNA

The shRNA target sequence for human HSF1 (NM_005526.4) was selected using the RNAi Target Sequence Selector (Clontech, Mountain View, CA, USA). The target sequences were: shHSF1 - 5’GCA GGT TGT TCA TAG TCA GAA-3’ (1994-2013 in NM_005526.4), shHSF1.2 - 5’CCT GAA GAG TGA AGA CAT A (526-544), and shHSF1.3 - 5’ CAG TGA CCA CTT GGA TGC TAT (1306-1326). The negative control sequence was 5’-ATG TAG ATA GGC GTT ACG ACT. Sense and antisense oligonucleotides were annealed and inserted into the pLVX-shRNA vector (Clontech) at *Bam*HI/*Eco*RI site. Infectious lentiviruses were generated by transfecting DNA into HEK293T cells and virus-containing supernatant was collected. Human MCF7 cells were transduced with lentiviruses following the manufacturer’s instructions and selected using a medium supplemented with 1 μg/ml puromycin (Life Technologies / Thermo Fisher Scientific, Waltham, MA, USA).

### HSF1 functional knockout using the CRISPR/Cas9 editing system

To remove the human *HSF1* gene, Edit-R Human HSF1 (3297) crRNA, Edit-R tracrRNA, and Edit-R hCMV-PuroR-Cas9 Expression Plasmid (Dharmacon, Lafayette, CO, USA) were introduced into MCF7 cells using DharmaFECT Duo (6 µg/ml) (Dharmacon) according to producer’s instruction. Transfected cells were enriched by puromycin (2 µg/ml) selection for 4 days. Afterward, single clones were obtained by limiting dilution on a 96-well plate. The efficiency of the HSF1 knockout was monitored by Western blot. Out of 81 tested clones, two individual clones with the *HSF1* knockout (KO#1 and KO#2) and six pooled control clones (MIX) were chosen for the next experiments. Among individually tested HSF1-targeting crRNAs, only two were effective (target sequences: GTGGTCCACATCGAGCAGGG and AAAGTGGTCCACATCGAGCA, both in exon 3 on the plus strand). For validation experiments, a new model was created: Edit-R Human HSF1 (3297) crRNAs (GGTGTCCGGGTCGCTCACGA in exon 1 on the minus strand and AAAGTGGTCCACATCGAGCA in exon 3 on the plus strand), Edit-R tracrRNA (Dharmacon), and eSpCas9-GFP protein (#ECAS9GFPPR, Merck KGaA) were introduced into MCF7 and T47D cells using Viromer® CRISPR (Lipocalyx GmbH, Halle (Saale), Germany) according to the manual provided by the producer. Single clones were obtained by limiting dilution on a 96-well plate. The efficiency of the HSF1 knockout was monitored by western blot and confirmed by sequencing (Genomed, Warszawa, Poland). Five (T47D) or six (MCF7) individual unaffected clones (HSF1+) or with the HSF1 functional knockout (HSF1−) were pooled each time before analyzes.

### Protein extraction and Western blotting

Whole-cell extracts were prepared using RIPA buffer supplemented with CompleteTM protease inhibitors cocktail (Roche) and phosphatase inhibitors PhosStopTM (Roche, Indianapolis, IN, USA). Proteins (20-30 μg) were separated on 10% SDS-PAGE gels and blotted to a 0.45-μm pore nitrocellulose filter (GE Healthcare, Europe GmbH, Freiburg, Germany) using Trans Blot Turbo system (Bio-Rad, Hercules, CA, USA) for 10 min. Primary antibodies against HSF1 (1:4,000, ADI-SPA-901, Enzo Life Sciences, Farmingdale, NY, USA), HSP90 (1:2,000, ADI-SPA-836, Enzo Life Sciences), anti-HSP70 (1:2,000, ADI-SPA-810, Enzo Life Sciences), anti-HSP105 (1:600, #3390-100, BioVision, Milpitas, CA, USA), ESR1 (1:2,000, #8644, Cell Signaling Technology, Danvers, MA, USA), and ACTB (1:25,000, #A3854, Merck KGaA) were used. The primary antibody was detected by an appropriate secondary antibody conjugated with horseradish peroxidase (Thermo Fisher Scientific, Waltham, MA, USA) and visualized by ECL kit (Thermo Fisher Scientific) or WesternBright Sirius kits (Advansta, Menlo Park, CA, USA). Imaging was performed on x-ray film or in a G:BOX chemiluminescence imaging system (Syngene, Frederick, MD, USA). The experiments were repeated in triplicate and blots were subjected to densitometric analyses using ImageJ software to calculate relative protein expression after normalization with loading controls (statistical significance of differences was calculated using T-test).

### Total and nascent RNA isolation, cDNA synthesis, and RT-qPCR

For nascent RNA labeling, 500 μM of 4-thiouridine (Cayman Chemical, Ann Arbor, MI, USA) was added to control and E2-treated cells for the duration of the treatment (4h). Next, total RNA was isolated using the Direct-ZolTM RNA MiniPrep Kit (Zymo Research, Irvine, CA, USA), digested with DNase I (Worthington Biochemical Corporation, Lakewood, NJ, USA), and cleaned with RNAClean XP beads (Beckman Coulter Life Science, Indianapolis, USA). Five micrograms of total RNA from each sample were taken for nascent RNA fraction isolation using methane tiosulfonate (MTS) chemistry according to (Duffy and Simon, 2016). After the biotinylation step using MTSEA-biotin-XX (Biotium, Fremont, CA, USA), s4U-RNA was cleaned with RNAClean XP beads and isolated using μMacs Streptavidin Kit (Miltenyi Biotec, Bergisch Gladbach, Germany) as described (Garibaldi et al., 2017). Total RNA (1 μg) and nascent RNA (isolated from 5 μg of total RNA) from each sample were converted into cDNA as described (Kus-Liśkiewicz et al., 2013). Quantitative PCR was performed using a BioRad C1000 Touch^TM^ thermocycler connected to the head CFX-96. Each reaction was performed at least in triplicates using PCR Master Mix SYBRGreen (A&A Biotechnology, Gdynia, Poland). Expression levels were normalized against *GAPDH*, *ACTB*, *HNRNPK*, *HPRT1*, if not stated otherwise. The set of delta-Cq replicates (Cq values for each sample normalized against the geometric mean of four reference genes) for control and tested samples were used for statistical tests and estimation of the p-value. Shown are median, maximum, and minimum values of a fold-change vs untreated control. The primers used in these assays are described in Table S1.

### Clonogenic assay

Cells were plated onto 6-well dishes (1 × 10^3^ cells per well) and cultured for 14 days. Afterward, cells were washed with the phosphate-buffered solution (PBS) and fixed with methanol. Colonies were stained with 0.2% crystal violet, washed, and air-dried. Colonies were counted manually.

### Aldefluor assay

The assay was performed using a kit from Stem Cell Technology (Vancouver, Canada, #01700) according to the protocol. Cells (6 x 10^5^) were harvested by trypsinization and resuspended in 1 ml Aldefluor Buffer. After the addition of 5 μl of BODIPY-aminoacetaldehyd (BAAA), the substrate for aldehyde dehydrogenase (ALDH), and a brief mixing, 500 μl of the cell suspension (3 × 10^5^) was immediately transferred to another tube supplemented with 5 μl of diethylaminobenzaldehyde (DEAB), a specific inhibitor of ALDH, and pipetted to mix evenly. Tubes were incubated at 5% CO_2_, 37 °C for 60 min. Cells were collected by centrifugation and resuspended in Aldefluor Buffer. Analyses were performed using the BD Canto III cytometer (Becton Dickinson, Franklin Lakes, NJ, USA).

### Proliferation test

Cells (2 ×10^4^ cells per well) were seeded and cultured in 12-well plates. At the indicated time, cells were washed with PBS, fixed in cold methanol, and rinsed with distilled water. Cells were stained with 0.1% crystal violet for 30 min, rinsed with distilled water extensively, and dried. Cell-associated dye was extracted with 1 ml of 10% acetic acid. Aliquots (200 μl) were transferred to a 96-well plate and the absorbance was measured at 595 nm (Synergy2 microtiter plate reader, BioTek Instruments). All values on day 0 were normalized to the optical density of wild-type cells (shown as 1.0). Grow curves are shown as the ratio of the absorbance on days 2, 4, and 6 against day 0 and were calculated from three to six independent experiments, each in 2-3 technical replicates. For each dataset the normality of distribution was assessed by the Shapiro-Wilk test and depending on data distribution homogeneity of variances was verified by the Levene test or Brown-Forsythe test. For analysis of differences between compared groups with normal distribution, the quality of mean values was verified by the ANOVA test with a pairwise comparison done with the HSD Tukey test or Games-Howell test and Tamhane test depending on the homogeneity of variance. In the case of non-Gaussian distribution, the Kruskal–Wallis ANOVA was applied for the verification of the hypothesis on the equality of medians with Conover-Iman’s test for pairwise comparisons.

### Proximity Ligation Assay

To detect the ESR1/HSP90 and ESR1/HSF1 interactions, the DuoLink in situ Proximity Ligation Assay (PLA) (Merck KGaA) was used according to the manufacturer’s protocol. Cells were plated onto Nunc® Lab-Tek® II chambered coverglass (#155383, Nalge Nunc International, Rochester, NY, USA) one day before the experiment. Cells were fixed for 15 minutes with 4% paraformaldehyde solution in PBS, washed in PBS, and treated with 0.1% Triton-X100 in PBS for 5 minutes. After washing, slides were incubated in Blocking Solution and immunolabeled (overnight, 4 °C) with primary antibodies diluted in the DuoLink® Antibody Diluent: rabbit anti-HSP90 (1:200; #ADI-SPA-836, Enzo Life Science) or rabbit anti-HSF1 (1:300; ADI-SPA-901, Enzo Life Sciences) and mouse anti-ERalpha (1:200; # C15100066, Diagenode, Liège, Belgium); negative controls were proceeded without one primary antibody or both. Then the secondary antibodies with attached PLA probes were used. Signals of analyzed complexes were observed using Carl Zeiss LSM 710 confocal microscope with ZEN navigation software; red fluorescence signal indicated proximity (< 40 nm) of proteins recognized by both antibodies (Fredriksson et al., 2002). Z-stacks images (12 slices; 5.5 μm) were taken at ×630 magnification. From each experimental condition, spots from 10 images were identified using Photoshop (*Red Channel → Select → Color Range*) and counted (*Picture → Analysis → Record the measurements*). Next, the mean number of spots per cell (nucleus, cytoplasm) in each image was calculated. Experiments were repeated three times. Outliers were determined using the Grubbs, Tuckey criterion, and QQ plot. For each dataset, the normality of distribution was assessed by the Shapiro-Wilk test. In the case of non-Gaussian distribution, the Kruskal–Wallis ANOVA was applied for the verification of the hypothesis on the equality of medians with Conover-Iman’s test for pairwise comparisons.

### Global Gene Expression Profiling

Total RNA was isolated from all MCF7 cell variants (untreated, treated with 10 nM E2 for 4 h, conditions based on (Vydra et al. 2019)) using the Direct-ZolTM RNA MiniPrep Kit (Zymo Research) and digested with DNase I (Worthington Biochemical Corporation). For each experimental point, RNAs from three biological replicates were first tested by RT-qPCR for the efficiency of treatments, then pooled. cDNA libraries were sequenced by Illumina HighSeq 1500 (run type: paired-end, read length: 2 × 76 bp). Raw RNA-seq reads were aligned to the human genome hg38 in a Bash environment using tophat2 (Kim et al., 2013) with Ensembl genes transcriptome reference. Aligned files were processed using Samtools (Li et al., 2009). Furthermore, reads aligned in the coding regions of the genome were counted using FeatureCounts (Liao et al., 2014). Further data analyses were carried out using the R software package v. 3.6.2 (R Foundation for Statistical Computing; http://www.r-project.org). Read counts were normalized using DESeq2 (Lowe et al., 2014), then normalized expression values were subjected to differential analysis using NOISeq package v. 3.12 (Tarazona et al., 2015) (E2 versus Ctr in all cell variants separately). To find common genes between samples, lists of differentiating genes were compared and Venn diagrams were performed (package VennDiagram v 1.6.20 from CRAN). Heatmaps of normalized read counts or log2 fold changes (E2 versus Ctr) for genes shared between samples were generated (package pheatmap v. 1.0.12 from CRAN). The hierarchical clustering of genes was based on Euclidean distance. Colors are scaled per row. Geneset enrichment analysis was performed as follows: From the count matrices, we filtered out all the genes with less than 10 reads in each of the libraries. Then, we analyzed the gene-level effects of E2 stimulation of cells with normal/decreased HSF1 levels, performing the DESeq2 test for paired samples, with pairs defined by the cell type (3 cell types with normal- HSF1 level: WT, SRC, MIX, and 3 cell types with decreased HSF1 level: shHSF1, KO#1 and KO#2). Finally, we performed the geneset enrichment analysis in the same way as for the TCGA data (see below for details) - for each test, genes were ranked according to their Minimum Significant Difference, CERNO test from tmod package was used to find enriched terms, tmodPanelPlot function was used to visualize the results. The raw RNA-seq data were deposited in the NCBI GEO database; acc. no. GSE159802.

### Chromatin Immunoprecipitation and ChIP-qPCR

The ChIP assay was performed according to the protocol from the iDeal ChIP-seq Kit for Transcription Factors (Diagenode) as described in detail in (Vydra et al., 2019). For each IP reaction, 30 µg of chromatin and 4 μl of mouse anti-ERalpha monoclonal antibody (C15100066, Diagenode) was used. For negative controls, chromatin samples were processed without antibody (mock-IP). Obtained DNA fragments were used for global profiling of chromatin binding sites or gene-specific ChIP-qPCR analysis using specific primers covering the known estrogen response elements (ERE). The set of delta-Cq replicates (difference of Cq values for each ChIP-ed sample and corresponding input DNA) for control and test sample were used for ESR1 binding calculation (as a percent of input DNA) and estimation of the p-values. ERE motifs in individual peaks were identified using MAST software from the MEME Suite package (Bailey, 2011). The sequences of used primers are presented in Table S2.

### Global Profiling of Chromatin Binding Sites

In each experimental point, four ChIP biological replicates (each from 30 μg of input chromatin) were collected and combined in one sample before DNA sequencing. Immunoprecipitated DNA fragments and input DNA were sequenced using the Illumina HiSeq 1500 system and QIAseq Ultralow Input Library Kit (run type: single read, read length: 1 × 65 bp). Raw sequencing reads were analyzed according to standards of ChIP-seq data analysis as described below. Quality control of reads was performed with FastQC software (www.bioinformatics.babraham.ac.uk/projects/fastqc) and low-quality sequences (average phred < 30) were filtered out. Remained reads were aligned to the reference human genome sequence (hg19) using the Bowtie2.2.9 program (Langmead and Salzberg, 2012). Individual peaks (Ab-ChIPed samples versus input DNA) and differential peaks (17β-estradiol-treated versus untreated cells) were detected using MACS software (Feng et al., 2012), whereas the outcome was annotated with Homer package (Heinz et al., 2010). Peak intersections and their genomic coordinates were found using Bedtools software (Quinlan and Hall, 2010). The input DNA was used as a reference because no sequences were obtained using a mock-IP probe. The locations of identified ESR1-binding sites were compared to genomic coordinates of E2-induced HSF1 peaks from our previous ChIP-seq analysis (NCBI GEO database; acc. no. GSE137558). We defined ESR1/HSF1 binding sites as “common” if at least the center of one peak was within the corresponding peak. Dot plots showing peak size distribution were generated using MedCalc Statistical Software version 19.2.1 (MedCalc Software Ltd, Ostend, Belgium; https://www.medcalc.org; 2020). ChIP-Seq heatmaps were prepared using peakHeatmap function from ChIPseeker Bioconductor package (version 1.26.2), with margins of 3,000 nucleotides upstream and downstream from the promoter. The raw ChIP-seq data were deposited in the NCBI GEO database; acc. no. GSE159724.

### Chromosome conformation capture assay (3C)

The procedure was carried out according to the protocol from (Deng and Blobel, 2017). In brief, 1 × 10^7^ cells per sample were trypsinized and fixed with 1% formaldehyde in 1x PBS. Crosslinking was quenched by 0.125 M glycine and cells were lysed (10 mM Tris pH 8.0, 10 mM NaCl, 0.2% NP-40, protease inhibitors). Cell nuclei were resuspended in *HindIII* RE buffer (10 mM Tris pH 8.0, 50 mM NaCl, 10 mM MgCl_2_, 100 μg/ml BSA) and incubated sequentially with 0.3% SDS (1.5 hours) and 1.8% Triton X-100 (1.5 hours) at 37 °C with rotation. Chromatin was cleaved using 450U *HindIII* restriction enzyme (BioLabs, Massachusetts, USA) at 37 °C overnight and diluted 15-fold in ligation buffer (50 mM Tris pH 7.5, 10 mM MgCl_2_, 10 mM DTT, 1% Triton X-100, 100 μg/ml BSA). Ligation was carried out using 4,000U T4-DNA ligase (EURx) at 16°C overnight, in the presence of 1 mM ATP. All samples were de-crosslinked (65 °C, overnight with mixing), RNase A and Proteinase K treated, and DNA was isolated using standard Phenol/Chloroform/Isoamyl alcohol purification method. Precipitated DNA was dissolved in 10 mM Tris pH 8.0 and used as a template in PCR analyses. The primers used are listed in Table S3.

**Analysis of human patient TCGA data** (performed using R version 3.6.2.).

*<u>Data retrieval</u>*. Clinical and RNA-seq (HTSeq counts) data from the TCGA (The Cancer Genome Atlas) breast cancer (BRCA) project was downloaded (1102 total samples) and prepared using the TCGAbiolinks package (version 2.14) (Colaprico et al., 2016). An additional file with clinical data containing ER receptor status, ‘nationwidechildrens.org_clinical_patient_brca.txt’, was downloaded directly from the GDC repository https://portal.gdc.cancer.gov. Molecular subtype classification (according to (Berger et al., 2018)) was retrieved through TCGAbiolinks.

*<u>Cases selection.</u>* Counts were log CPM normalized with the *cpm* function from the edgeR package (version 3.28.1) (Robinson et al., 2010). Then we selected four groups (called A-D) of patients: ER-positive/negative with high/low HSF1 expression level using the following clinical and expression (log CPM) criteria: ER-positive if er_status_by_ihc: “Positive”, er_status_ihc_Percent_Positive: “90-99%” and expression level of *ESR1* > 6, ER-negative if er_status_by_ihc: “Negative” and expression level of *ESR1* < 6, HSF1-high (low) if the expression of *HSF1* was above 67 (below 33) percentile across all TCGA_BRCA cases. We also excluded cases classified to HER2-enriched and basal-like subtypes from the ER-positive group and luminal A (LumA), luminal B (LumB), and normal-like subtypes from the ER-negative group. In reduced groups (called a-d) only the luminal A and basal-like cases were analyzed.

*<u>Survival analysis</u>* was performed using the *survfit* function from the *survival* package (version 3.1-8) and plotted with the *ggsurvplot* function from the *survminer* package (version 0.4.6) (Therneau and Grambsch, 2000).

*<u>MDS plots</u>* were used to visualize differences between patients. We performed multidimensional scaling (MDS) with *MDSplot* function from edgeR and plotted the results with ggplot2 (version 3.3.2) (Wickham, 2016).

*<u>Differential expression analysis</u>* was done with the edgeR package (Robinson et al., 2010). Lowly expressed genes were filtered out by filterByExpr function with default parameters, resulting in 24,696 genes kept for statistical analysis. Then we performed a quasi-likelihood F test for all groups’ combinations one-vs-one and two obvious two-vs-two cases: ER+ versus ER− and HSF1-high versus HSF1-low (using mean expression levels for joined groups, with the weight of 0.5). P-values were corrected for multiple testing using the Benjamini and Hochberg method.

*<u>Comparison of the number of genes differentially expressed between groups.</u>* To compare the size of differences identified in each test we plotted the cumulative distributions of FDR (False Discovery Rate). Each test was repeated 10 times with the groups randomly subsampled to equal size of 30 (or 28 in case of reduced groups) to avoid the p-values being affected by the group size inequality. Results were averaged and plotted with ggplot2 (Wickham, 2016).

*<u>Geneset enrichment analysis</u>.* For gene set enrichment analysis we selected hallmark, BioCarta, Reactome, and PID genesets from MSigDB (v7.0) (Subramanian et al., 2005) and merged it with the list of pathways downloaded from KEGG. DESeq2 was used to calculate log-fold changes with its standard error (Love et al., 2014). Then all genes were ordered according to their Minimum Significant Difference (MSD) calculated as |logFC| - 2\**logFC_standard_error* and tested for enrichment using the CERNO test (Zyla et al., 2019) from the tmod package (version 0.44) (Weiner 3rd and Domaszewska, 2016). The most significant results (effect size > 0.65, p-value < 0.001 at least in one comparison) and results for genesets related to the biological processes of interest were visualized with the *tmodPanelPlot* function.

## Acknowledgments

The authors thank Mrs. Krystyna Klyszcz and Urszula Bojko for technical assistance.

## Author Contributions

conceptualization, N.V., P.J., and W.W.; methodology, N.V., P.J., P.K., K.M., A.T-J., and W.W.; validation, N.V., P.J., and A.T-J.; formal analysis, N.V., P.J., P.K., T.S., A.T-J., A.J.C., A.K., and R.J.; data curation, T.S. and P.J.; investigation, N.V., P.J., K.M., A.T-J., B.W., B.G., and W.W.; writing – original draft preparation, N.V., P.J., P.K., and W.W.; writing – review and editing, N.V., P.J., and W.W.; visualization, N.V., P.J., P.K., and W.W.; supervision, N.V., P.J., R.J., M.K., and W.W.; project administration, N.V., and W.W.; funding acquisition, N.V., M.K., and W.W.

## Supplementary Files: Supplementary Figure legends

**Fig. S1.**
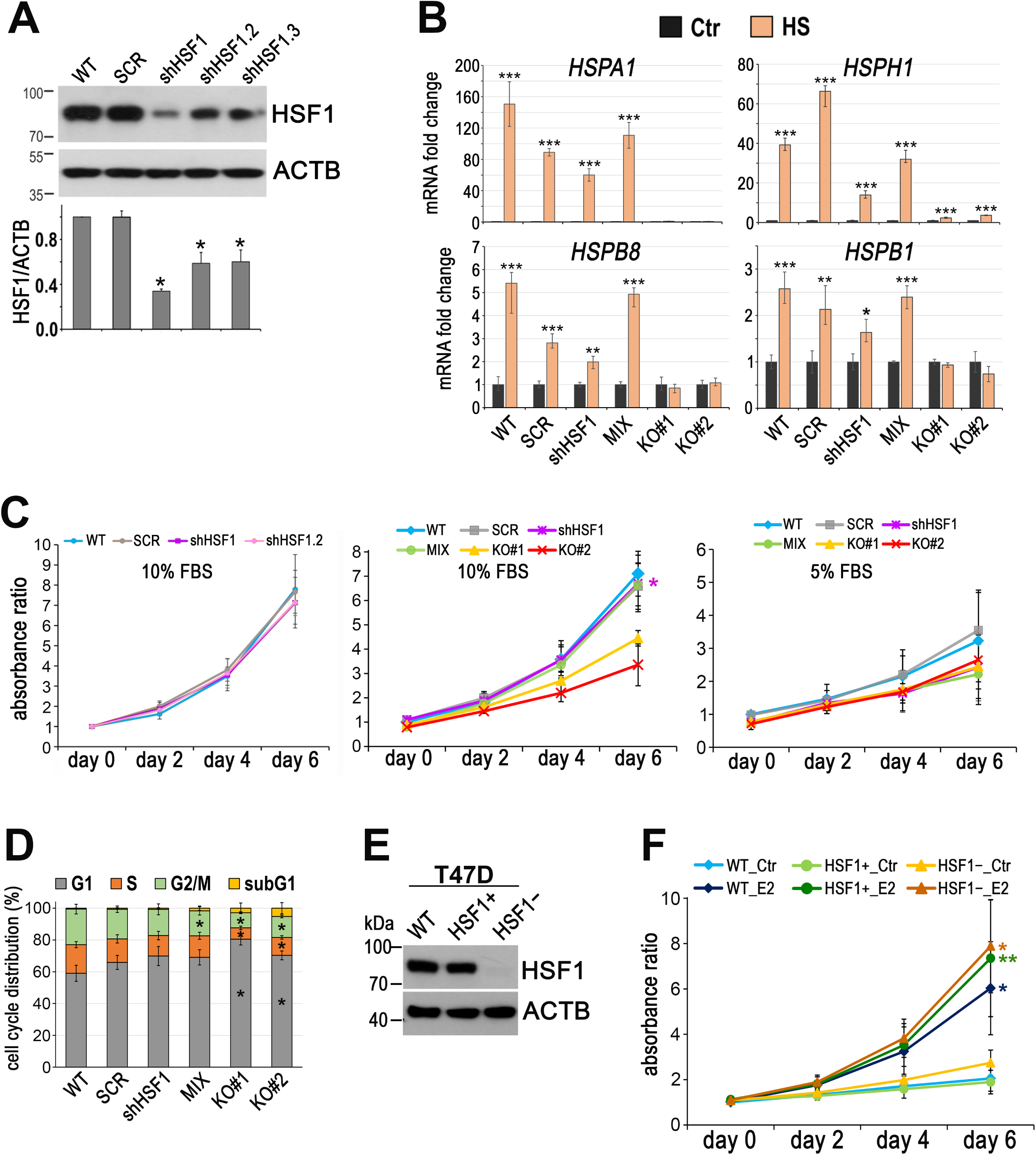
Characterization of HSF1-deficient cell variants. (**A**) Western blot analysis of HSF1 level in unmodified MCF7 cells (WT) and variants stably transduced with non-specific shRNA (SCR) or with three different HSF1-specific shRNAs (shHSF1). Actin (ACTB) was used as a protein loading control. The graph below shows the results of densitometric analyses of HSF1 immunodetection (n=3). *p < 0.05. (**B**) Expression of indicated *HSP* genes analyzed by RT-qPCR in MCF7 cell variants exposed to elevated temperature (HS: 43 °C/1h + recovery 37 °C/4h) in relation to untreated control (Ctr), ***p < 0.0001, **p < 0.001, *p < 0.05. (**C**) Cell growth curves under standard conditions (10% FBS) and 5% dextran-activated charcoal-stripped FBS assessed using crystal violet staining. Mean and standard deviation from three to four independent experiments (each in two technical replicates) are shown. *p < 0.05. (**D**) Cell cycle phases and sub-G1 distribution in sub-confluent cells at 72 hours after plating presented as 100% stacked column plots (mean±SD, n=3). *p < 0.05, significantly different to WT. (**E**) Western blot analysis of HSF1 level in T47D cell variants: unmodified (WT), and a combination of five control (HSF1+) and five HSF1-negative (HSF1−) clones arisen from single cells following CRISPR/Cas9 gene targeting. Actin (ACTB) was used as a protein loading control. (**F**) Cell growth curves of untreated (Ctr) and E2-treated T47D cell variants in phenol red-free media with 10% charcoal-stripped FBS assessed using crystal violet staining. Mean and standard deviation from three independent experiments (each in six technical replicates) are shown. **p < 0.001, *p < 0.05.

**Fig. S2.**
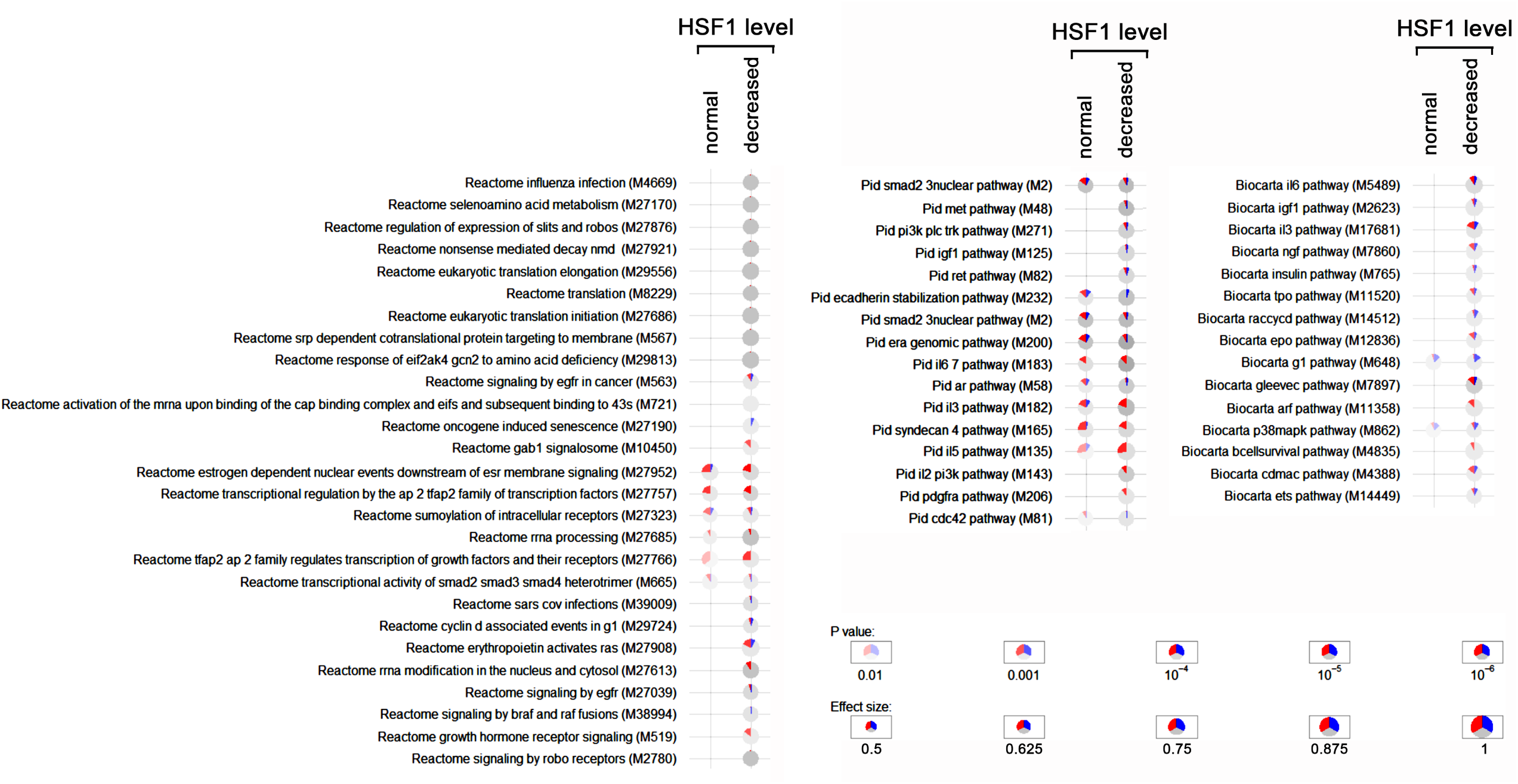
Geneset enrichment analysis showing differences between HSF1-proficient and HSF1-deficient MCF7 cells in response to E2 stimulation. Significant terms from the REACTOME, PID, and BIOCARTA subsets of the canonical pathways collection are shown. Blue – a fraction of down-regulated genes, red – fraction of up-regulated genes.

**Fig. S3.**
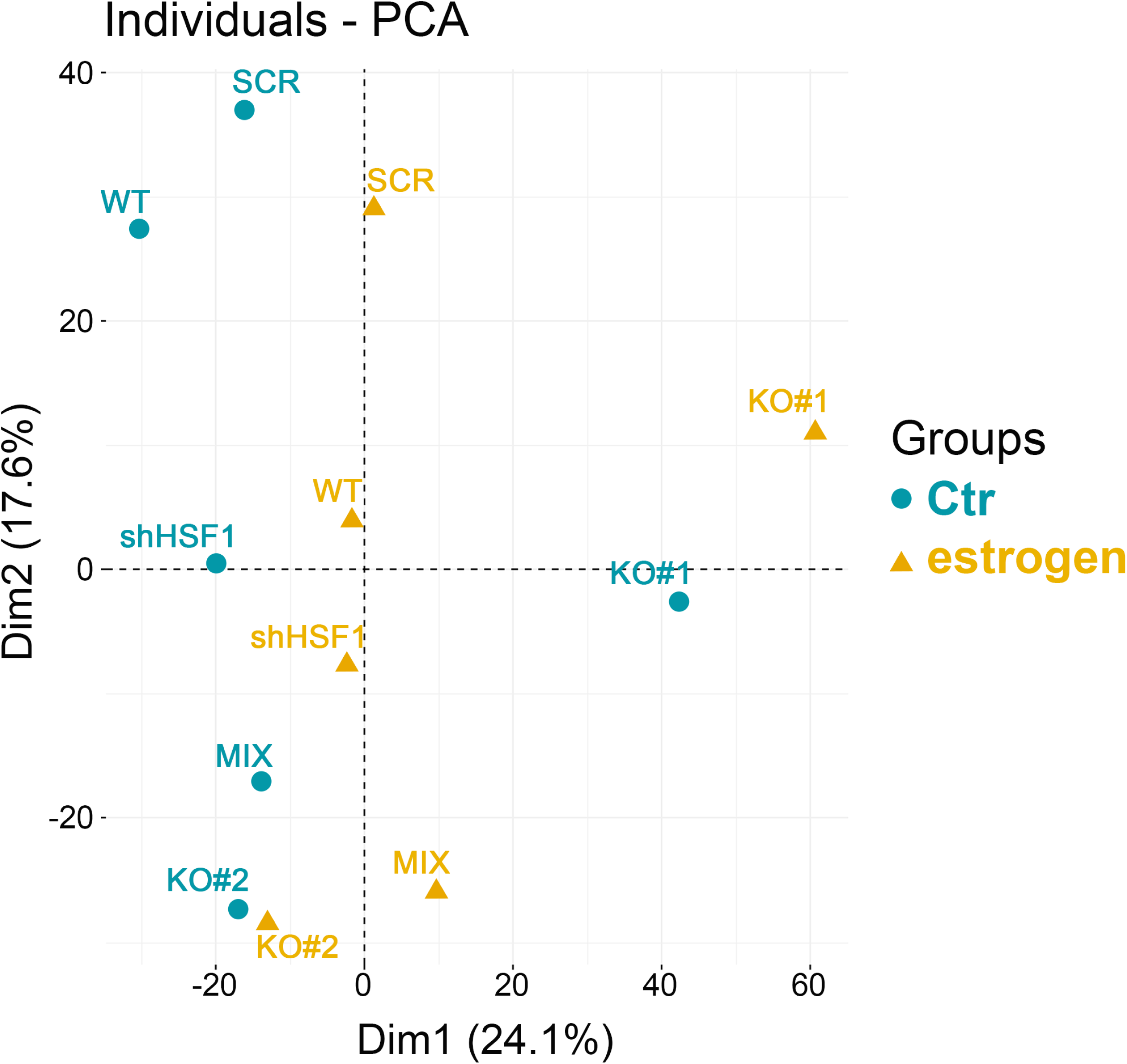
Principal component analysis (PCA) of normalized RNA-seq read counts. Gene expression changes were investigated in untreated (Ctr) and estrogen-treated (10 nM, 4 hours) MCF7 cell variants: unmodified (WT), stably transduced with non-specific shRNA (SCR), stably transduced with HSF1-specific shRNA (shHSF1); a combination of control clones arisen from single cells following CRISPR/Cas9 gene targeting (MIX), two HSF1 negative clones obtained by CRISPR/Cas9 gene targeting (KO#1, KO#2).

**Fig. S4.**
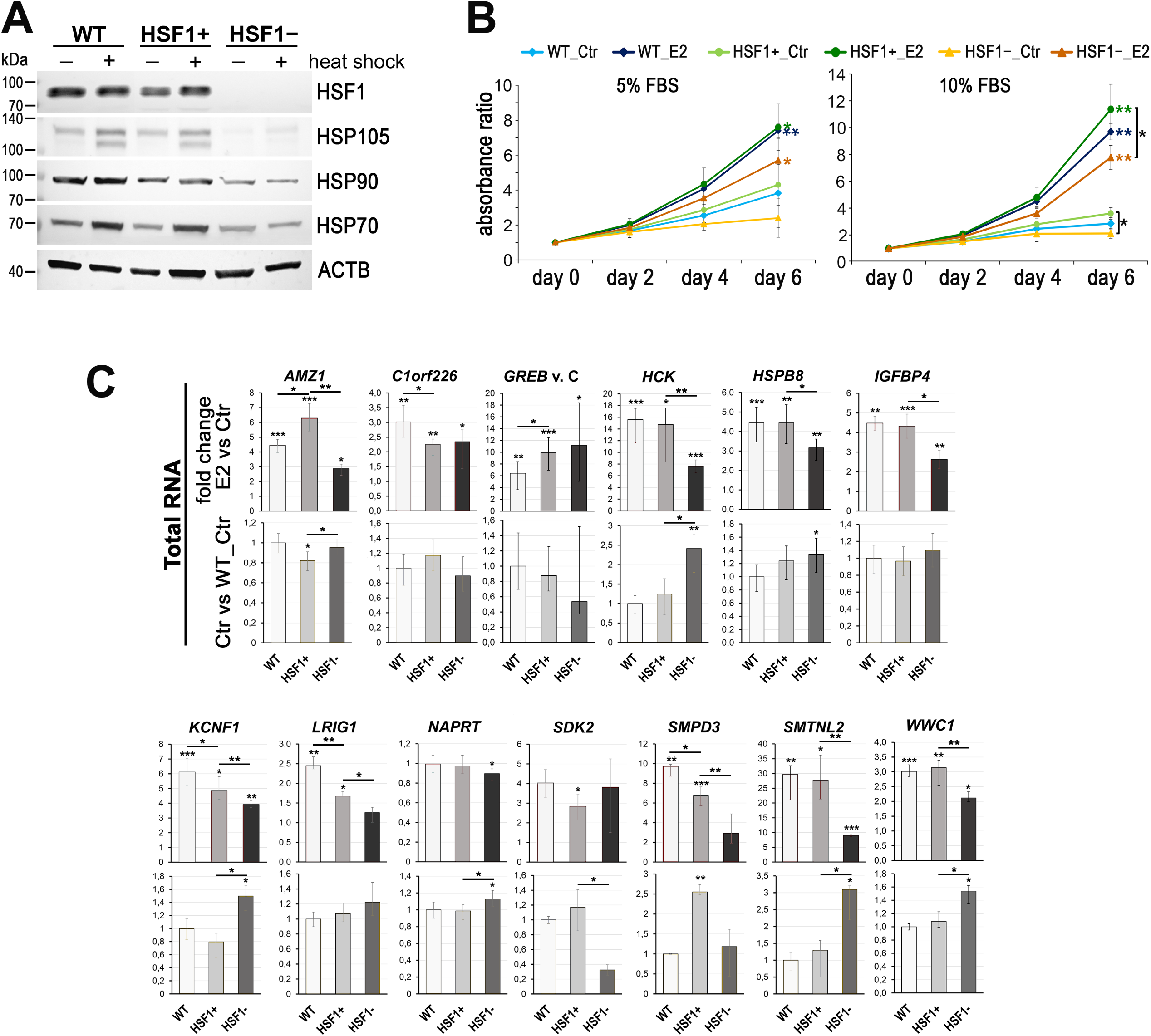
Characterization of MCF7 cell model created for validation using DNA-free CRISPR/Cas9 system. (**A**) Heat shock response assessed by western blot in unmodified (WT), HSF1+ (six HSF1-positive clones), and HSF1− (six HSF1-negative clones) cells. Actin (ACTB) was used as a protein loading control. Heat shock: 43 °C/1h + recovery 37 °C/6h. (**B**) Cell growth curves of untreated (Ctr) and E2-treated cells in phenol red-free media with 5 or 10% charcoal-stripped FBS assessed using crystal violet staining. Mean and standard deviation from three independent experiments (each in six technical replicates) are shown. **p < 0.001, *p < 0.05 (next to the curve – compared to the corresponding control, between curves – between cell variants). (**C**) Gene expression analyses by RT-qPCR using total RNA. The upper panel shows E2-stimulated changes, lower panel shows differences between untreated WT, HSF1+, and HSF1− cells. Ctr, untreated cells; E2, 17β-estradiol treatment (10 nM, 4 h). ***p < 0.0001, **p < 0.001, *p < 0.05 (above the bar – compared to the corresponding control, between the bars – between cell variants).

**Fig. S5.**
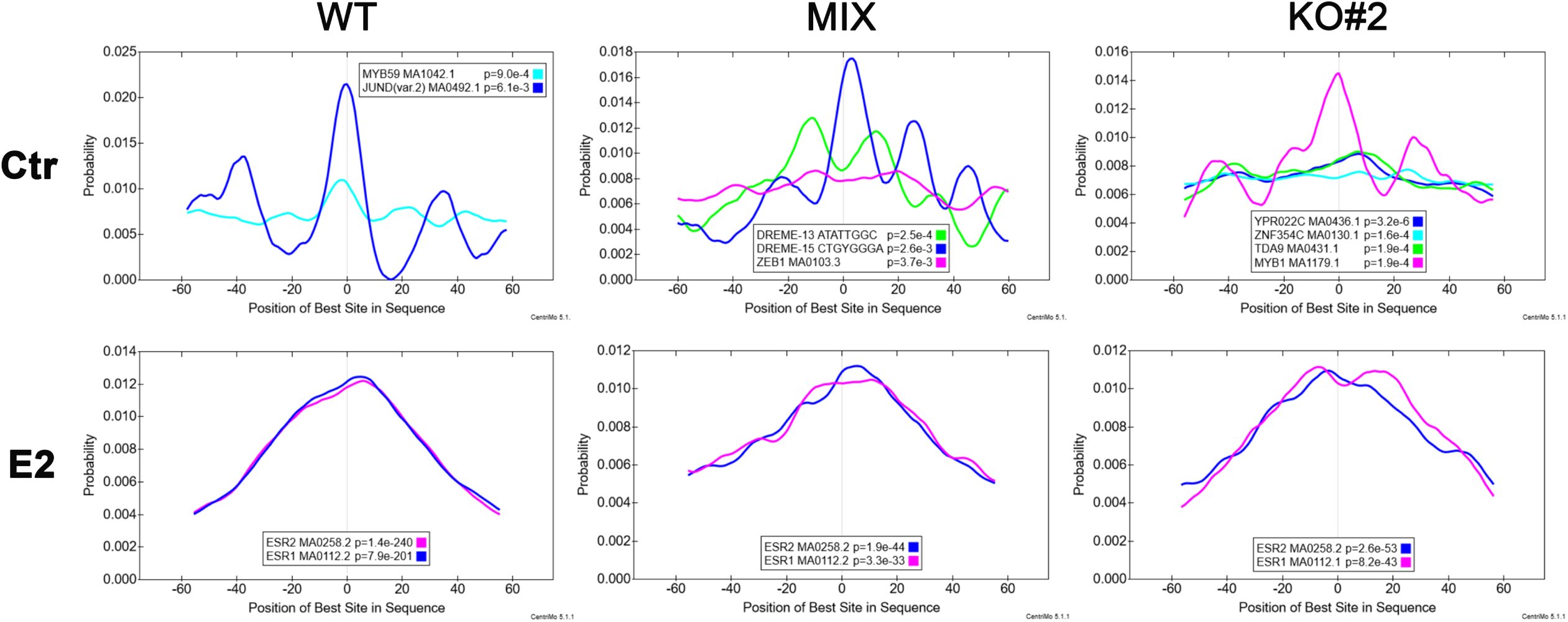
Top enriched motifs in ESR1 ChIP-seq peak regions. The CentriMo plots show the distribution of the given motifs in peaks from untreated (Ctr) and estrogen (E2)-treated (10 nM for 30 minutes) MCF7 cell variants. JASPAR motif names, IDs, and the p-value of the motif’s central enrichment in peaks are shown in the legend of each plot. MCF7 cell variants: wild type (WT), a combination of control clones arisen from single cells following CRISPR/Cas9 gene targeting (MIX), HSF1 negative clone obtained by CRISPR/Cas9 gene targeting (KO#2).

**Fig. S6.**
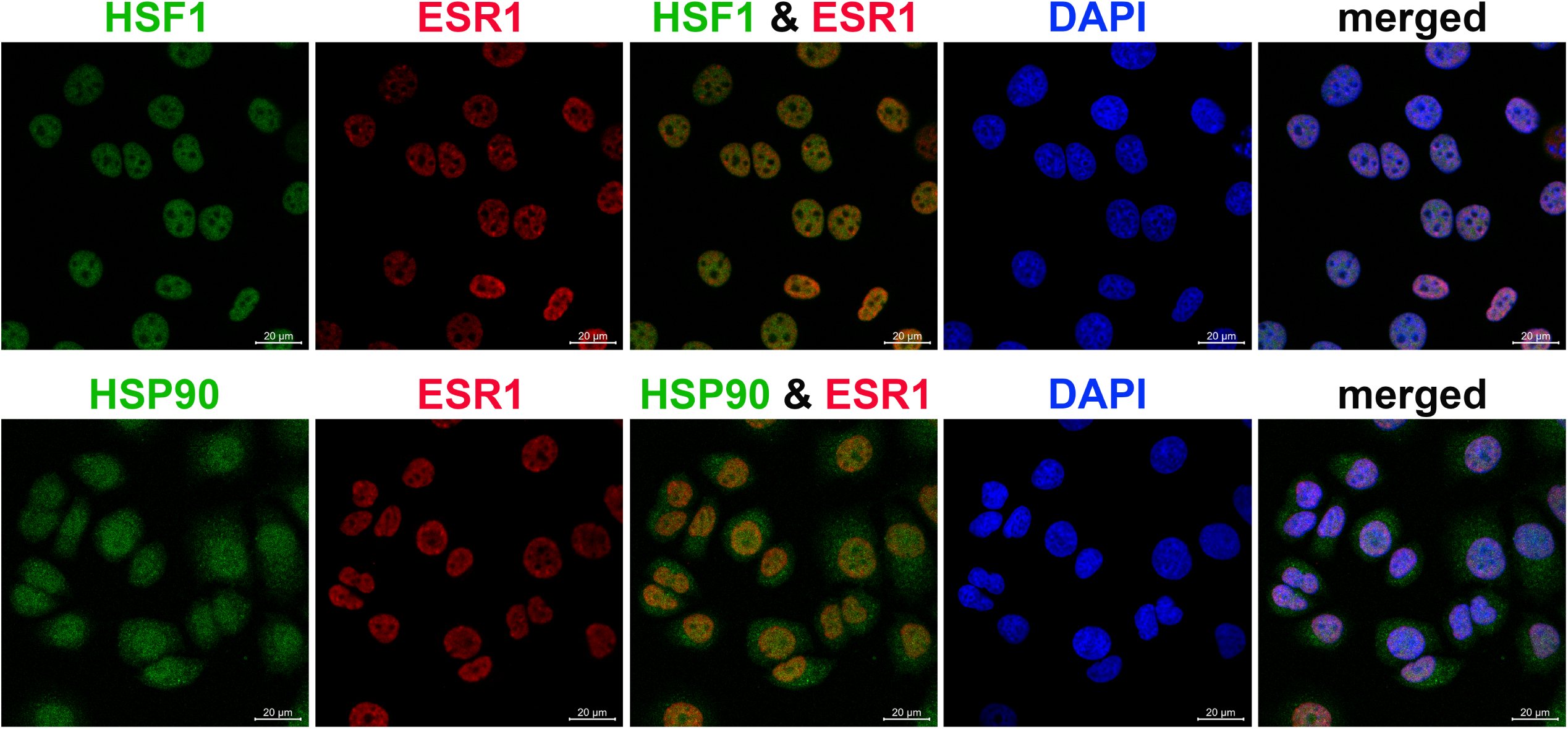
HSF1, HSP90, and ESR1 localization assessed by immunofluorescence in MCF7 cells. DNA was stained with DAPI. Scale bar, 20 μm.

**Fig. S7.**
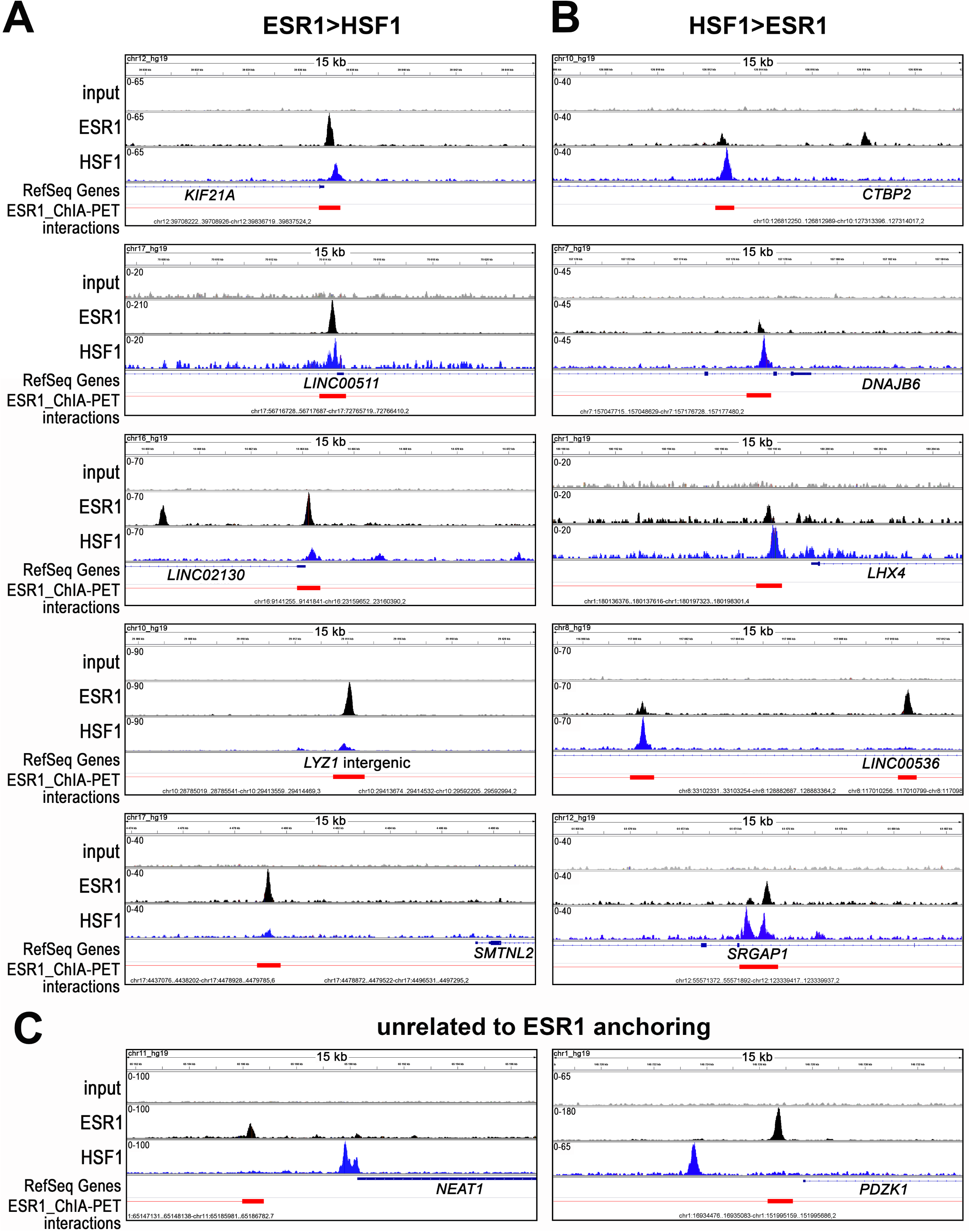
Examples of different patterns of ESR1 and HSF1 binding to chromatin. (**A**) Stronger binding of ESR1 than HSF1. (**B**) Stronger binding of HSF1 than ESR1. (**C**) HSF1 binding unrelated to ESR1 anchoring. Peaks identified by MACS in ChIP-seq analyses in wild-type MCF7 cells after E2 treatment (10 nM, 60 min) and corresponding ChIA-PET interactions (Fullwood et al., 2009) downloaded from ENCODE database and visualized by the IGV browser. The red bar shows the ESR1 anchor region (interacting loci), red line – a loop (the intermediate genomic span between the two anchors). The scale for each sample is shown in the left corner.

**Fig. S8.**
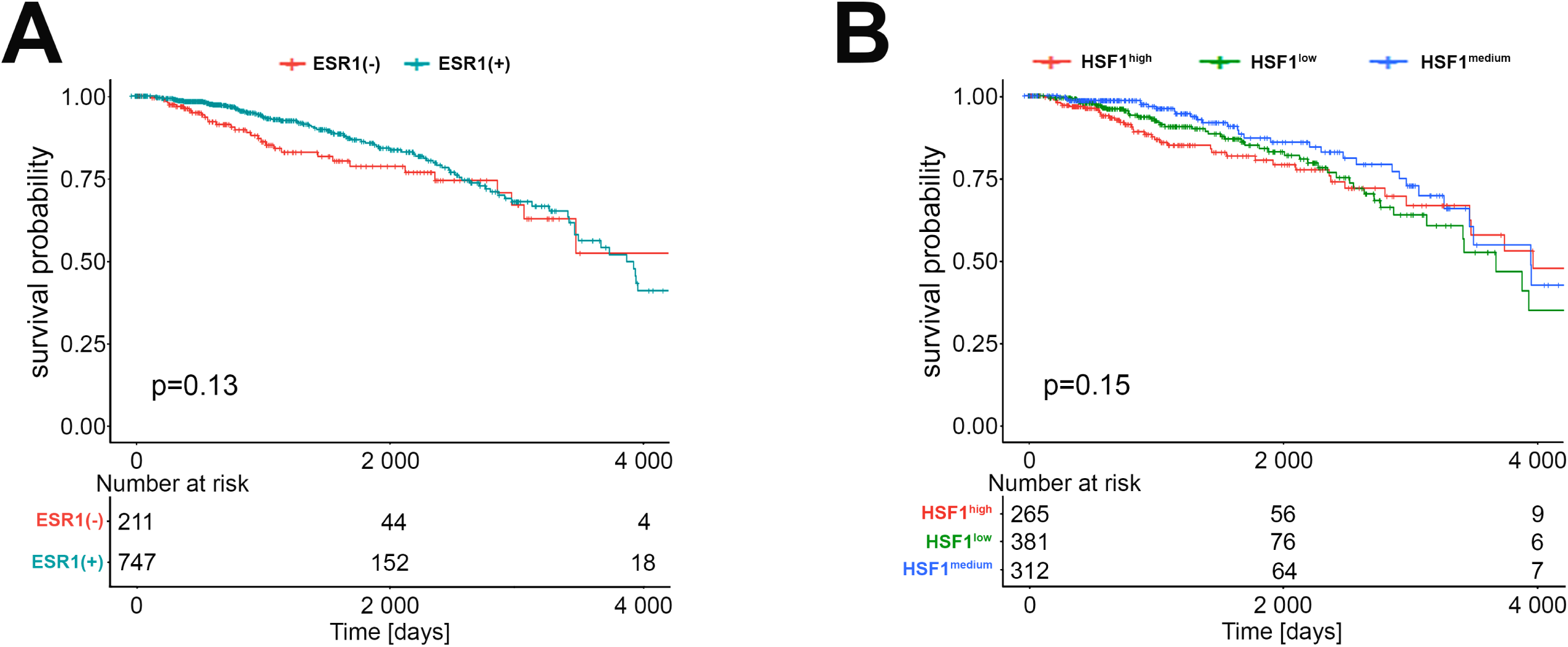
Effect of *ESR1* (**A**) and *HSF1* (**B**) transcript levels on survival in TCGA breast cancer patients analyzed using The Kaplan Meier Plotter.

**Fig. S9.**
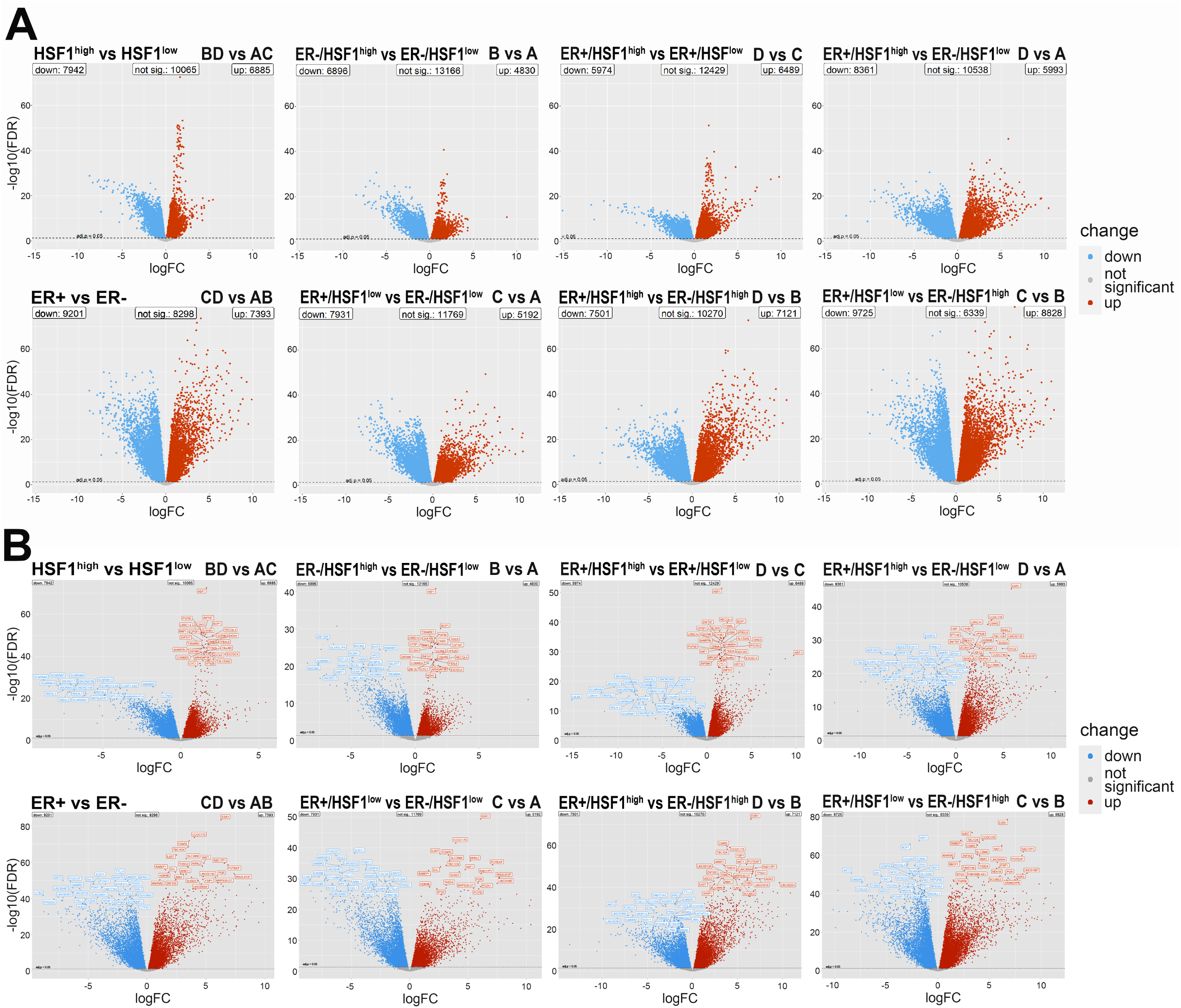
(**A**) Volcano plots visualizing differential expression patterns between two distinct groups of breast cancers with different levels of ESR1 and HSF1 expression (the same scale is kept). (**B**) Volcano plots with gene labels. The red points in the plots represent genes with statistically significant increased expression in the first group, the blue points - in the second group in the given comparison (adjusted p < 0.05).

**Fig. S10.**
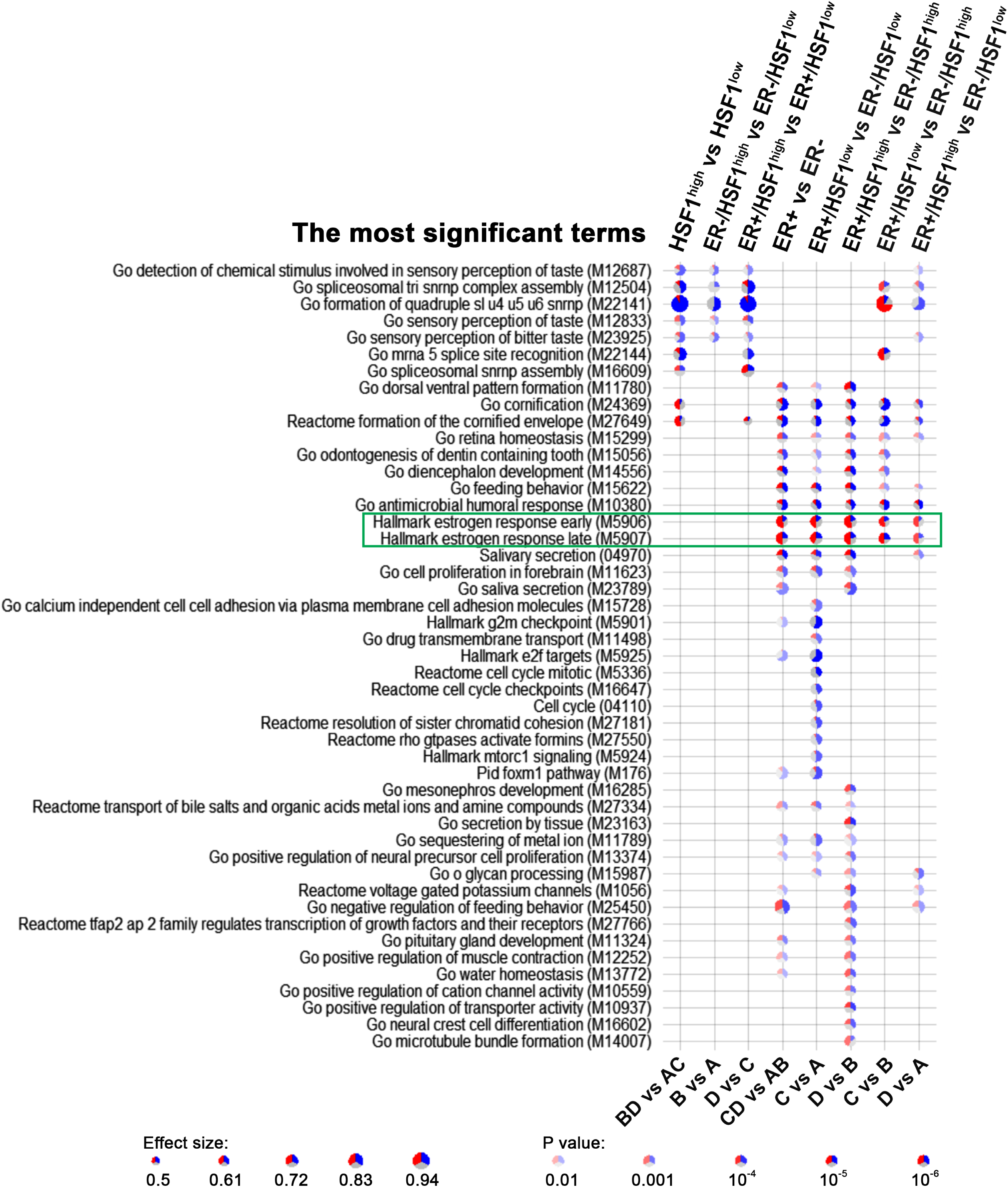
Geneset enrichment analyses showing the most significant terms differentiating ER+ and ER– breast cancers with different HSF1 levels. (in comparisons between groups selected in Fig. 6B). Blue – down-regulated genes, red – up-regulated genes. Terms related to estrogen response are marked with the green rectangle.

**Supplementary Dataset 1**. Summary table of RNA-seq results (normalized signals and expression fold changes after E2 treatment) in MCF7 cell variants with different levels of HSF1. xlsx document.

**Supplementary Dataset 2**. Summary tables of ChIP-seq results: characteristics of ESR1 binding in wild-type, HSF1-proficient (MIX), and HSF1-deficient (KO#2) MCF7 cells, untreated (Ctr) and after E2-stimulation. xlsx document.

**Supplementary Dataset 3**. Summary tables of ChIP-seq results: characteristics of ESR1 and HSF1 common binding regions in wild-type MCF7 cells, untreated (Ctr) and after E2-stimulation. xlsx document.

**Supplementary Dataset 4**. Differential expression tests between selected groups of breast cancer patients with different ESR1 and HSF1 statuses based on RNA-seq data deposited in the TCGA database. xlsx document.

**Supplementary methods:** Cell-cycle distribution. MEME-ChIP analyses. Immunofluorescence (IF).

## Supplementary Tables

**Table S1.** RT-qPCR primers for gene expression analyses.

**Table S2.** ChIP-qPCR primers for ESR1 binding analyses.

**Table S3.** PCR primers for chromosome conformation capture assay.

## Source Data

**Figs 1A, 4B, 5E - source data** (unedited gels and blots): the original files of the full raw unedited blots and gels and figures with the uncropped blots and gels with the relevant bands labelled.

## Supplementary methods

### Cell-cycle distribution

Cells (3 × 10^5^ per well) were plated onto 6-well plates. The next day medium was replaced and cells were grown for an additional 48 hours. Afterward, cells were harvested by trypsinization, rinsed with PBS, fixed with ice-cold 70% ethanol at -20 °C overnight. Cells were collected by centrifugation, resuspended in PBS containing RNase A (100 µg/ml), and stained with 100 µg/ml propidium iodide solution. DNA content was analyzed using flow cytometry to monitor the cell cycle changes.

### MEME-ChIP analyses

The consensus DNA sequences for ESR1 binding were identified *in silico* by Motif Analysis of Large Nucleotide Datasets (MEME-ChIP, version 5.1.1) (Bailey et al., 2009) using a 150-bp region centered on the summit point and visualized by CentriMo (Local Motif Enrichment Analysis) (Bailey and Machanick, 2012).

### Immunofluorescence (IF)

Cells were plated onto Nunc® Lab-Tek® II chambered coverglass (#155383, Nalge Nunc International, Rochester, NY, USA) and fixed for 15 minutes with 4% PFA paraformaldehyde solution in PBS, washed, treated with 0.1% Triton-X100 in PBS for 5 minutes, and washed again in PBS (3 x 5 minutes). IF imaging was performed using primary antibodies: anti-HSP90 (1:200; ADI-SPA-836, Enzo Life Science), anti-HSF1 (1:300; ADI-SPA-901, Enzo Life Sciences) or anti-ESR1 (1:200; C15100066, Diagenode) and secondary Alexa Fluor (488 or 594) conjugated antibodies (Abcam). Finally, the DNA was stained with DAPI. Images were taken using Carl Zeiss LSM 710 confocal microscope with ZEN navigation software.

